# Languages of the brain: fMRI dissection of the amodal networks for language, mathematics, and social knowledge

**DOI:** 10.64898/2025.12.19.695075

**Authors:** Antonio Moreno, Marie Amalric, Manon Pietrantoni, Séverine Becuwe, Bertrand Thirion, Christophe Pallier, Ghislaine Dehaene-Lambertz, Stanislas Dehaene

**Affiliations:** Cognitive Neuroimaging Unit, INSERM, CEA, CNRS, Université Paris Saclay, NeuroSpin center, 91191 Gif/Yvette, France; Collège de France, Université Paris Sciences Lettres (PSL), 11 Place Marcelin Berthelot, 75005 Paris, France; Inria, CEA, Université Paris Saclay

**Keywords:** Language, syntax, semantics, mathematics, arithmetic, number, geometry, Broca’s area, cerebellum, MRI, adolescence

## Abstract

The ability to compose complex mental representations by recombining simpler primitives is a characteristic of the human brain that manifests itself in a variety of domains such as spoken or written language, mathematics, or social reasoning. While it is known that these domains rest on partially distinct brain networks, they are rarely investigated together in the same paradigm. Here, we present a systematic functional MRI study of written and spoken word and sentence processing in each of these domains. In a true-false judgment task, adult participants were presented with a hierarchy of meaningless and meaningful stimuli bearing on various domains of general semantic knowledge and mathematics. The results show that distinct brain regions are activated by (1) syntax and semantics; (2) math versus non-math knowledge; (3) different domains of mathematics, such as geometry versus arithmetic; (4) social knowledge versus other knowledge. All of these regions are activated essentially identically in the written and spoken modalities. The results were replicated in a group of adolescents. Our experiments show that the human brain comprises distinct amodal networks for various domains of linguistic and semantic knowledge, and provide a simple paradigm to dissect them within a short fMRI session.

## Introduction

The ability to understand and produce spoken language is a striking characteristic of the human species that relies on an increasingly well-identified network of brain areas (Fedorenko et al., 2024). However, humans are also unique in the animal world in producing many other languages such as sign language, computer languages, music, or mathematics. The language-of-thought hypothesis stipulates that, in all of these domains, the singularity of the human brain lies in its ability to generate new symbols by recombining previous ones according to syntactic rules (Dehaene et al., 2022, 2025; Fodor, 1975; Frankland & Greene, 2020; Goodman et al., 2014; Quilty-Dunn et al., 2022). Even the ability to compose geometric drawings and signs may possess its own language-of-thought (Al Roumi et al., 2023; Dehaene et al., 2022; Sablé-Meyer et al., 2021).

At the computational level, it has been hypothesized that recursion may be the single common faculty that allows humans to expand their receptive and productive repertoires (Fitch, 2014; Hauser et al., 2002; Hauser & Watumull, 2017). At the neural level, however, much remains unknown about how, and in which brain circuits, this ability is implemented. Several authors initially postulated that a single brain circuit, possibly located in Broca’s area or the basal ganglia, underlies multiple domains of human competence such as language, music, and tool use (e.g. Fadiga et al., 2009; Fedorenko et al., 2009; Koelsch et al., 2002; Patel, 2003). However, with the advent of dedicated and higher-resolution brain-imaging studies, it became increasingly clear that dissociable brain circuits underlie domains that were once thought to rely on identical neural resources, such as language and music (Chen et al., 2021), language and logic (Fedorenko & Varley, 2016; Monti et al., 2009), language and mathematics (Amalric & Dehaene, 2016, 2017, 2019; Maruyama et al., 2012; Pinel et al., 2007; Varley et al., 2005), or natural versus computer languages (Liu et al., 2020; Liu & Bedny, 2025; but see Liu et al., 2024). Even within natural language, at least three different levels of composition can be distinguished, i.e. phonotactic, syntactic, and semantic composition, each of which thought to be associated with distinct parallel “unification” circuits linking different parts of prefrontal, temporal and inferior parietal cortex (Hagoort, 2013; Pallier et al., 2011). The language-of-thought hypothesis extends this idea by stipulating that similarly recursive systems are implemented in multiple parallel networks across both hemispheres of the human brain, with each network supporting a distinct cognitive domain—such as language, mathematics, music, theory-of-mind, etc. (Dehaene et al., 2022).

However, these domains are rarely investigated together at the brain level in the same participants, and most often only in pairs (e.g. Al Roumi et al., 2023; Amalric & Dehaene, 2016; Chen et al., 2021; Deen et al., 2015a; Fedorenko et al., 2011, 2012; Wang et al., 2019). Furthermore, even within the domain of language processing, it remains controversial whether language-related areas operate as a single system (Fedorenko et al., 2024; Shain et al., 2024) or can be dissociated along strict lines such as syntactic versus semantic composition (Hagoort, 2013; Pallier et al., 2011). In this context, the goal of the present study was to evaluate, within a single homogeneous paradigm, the brain regions involved in several different compositional systems of the human brain: syntactic word composition, elementary semantic composition, contextual embedding, social embedding (characteristic of theory-of-mind), and mathematics (including calculation, arithmetic principles, and geometry). We introduced a functional MRI paradigm that allowed these operations to be mapped within a short session of less than one hour. While previous research has often relied on extended textual stimuli, such as listening to an entire book such as The Little Prince (Pasquiou et al., 2023), a radio show (Deniz et al., 2019; Huth et al., 2016) or a short theory-of-mind story (Deen et al., 2015a; Saxe & Kanwisher, 2003), here we relied solely on single-sentence stimuli. All stimuli were approximately 3-seconds long, short enough to be presented in an event-related fMRI paradigm, with a simple verification task (is the sentence true or false?). Reducing mathematics or theory-of-mind questions to a single sentence, while controlling for length and word frequency, was challenging, yet we show that this minimal format still reliably activated distinct brain networks. This sentence paradigm also allowed us to systematically vary the modality of stimulus presentation.

Across subjects, all stimuli were presented either visually, in a rapid serial visual presentation, or auditorily, as short spoken sentences. Previous work has shown that many of the above brain networks respond amodally, with nearly identical activations to spoken and written materials across syntactic, semantic, social, and mathematical levels (Deen et al., 2015a; Dehaene et al., 2003; Deniz et al., 2019; Pinel et al., 2007). The present sentence paradigm extended this comparison to all tested domains, by systematically comparing visual and auditory presentation of the exact same sentences. We expected that all networks would exhibit modality-independent responses to spoken and written versions of the same content.

Here, we focused primarily on results obtained from 26 adults scanned at 3 Tesla. However, another major goal of our research was to evaluate whether the present design provides a stable and replicable paradigm that could eventually be used to assess the integrity of language, math and social knowledge networks in younger participants as in developmental patient groups with specific disorders such as specific language impairments, dyscalculia or autism spectrum disorders. To this end, we also collected data from a group of 15 adolescents tested with the same experimental paradigm and stimuli, and we briefly report how their results replicated those of the adult group.

Ongoing studies in our laboratory are evaluating variants of this paradigm in other populations, including congenitally blind adults and participants with high-functioning autism, using both standard 3 Tesla scans and ultra-high-resolution (1.2 mm) scans at 7 Tesla.

## Methods

### Participants

26 adult volunteers (24 males, 2 females, all right-handed, age range 19-45 years old, mean = 31.3 years old, standard deviation = 6.8) and 15 adolescent volunteers (all males, all right-handed, age range 14-15 years old, mean = 14.7 years old, standard deviation = 0.31) took part in the study. All were native speakers of French, had normal hearing and were fluent readers. The experiment was approved by the regional ethics committee, and all subjects gave written informed consent. Adults received 80 euros for their participation and teenagers, who were accompanied by one of their parents, received a small gift and a personalized diploma. Note that 10 of the adult participants belonged to the individual brain charting project (Pinho et al., 2018, 2020), for which they were repeatedly scanned several times with different paradigms. To remain consistent with the other acquisitions that they received, these participants were scanned using slightly different parameters. The 26 adult participants were pooled after verifying that their individual activations were similar.

### Stimuli

320 stimuli were created, each consisting in a short sequence of 6-13 French words (average 9.2 words), in spoken and written form. They were divided in 8 categories, each containing 40 stimuli in total. These categories were as follows:

● Lists of words: This category was used to present the same overall lexical items as in the other sentence conditions, while avoiding any possible parsing or semantic composition. To this end, half of these stimuli were lists of content words (nouns, adjectives, verbs), in a careful chosen pseudo-random order that prevented the formation of grammatical phrases from consecutive words, and the other half were lists of closed-class grammatical words (i.e. conjunctions, prepositions, determinants…). Some examples (as translated into English) are: “*Is the one while as without you for but in however”, or “Sweetness fish cinnamon grimace pole target sea”*.
● Semantically anomalous sentences (called “meaningless” for short): This category, inspired by Chomsky’s famous statement “Colorless green ideas sleep furiously”, contained sentences in which each content word had been replaced by another unrelated content word of the same lexical class (adjective, noun, etc). The aim was to obtain syntactically correct and easily parsed sentences, thus maximizing the activation of brain areas involved in syntactic composition, while reducing sentence-level meaning to a minimum. Some examples are “*The flag is an old analysis to cold manner” or “According to the wood, the feather is the choice of the seasonal discs”.* Obviously, the first few words might seem meaningful, but it quickly becomes clear that the successive words cannot be semantically integrated. Thus, we refer to this condition as “meaningless” for short, referring to the final status of the whole sentence, but acknowledging that they could elicit transient semantic integration efforts.
● Factual generic semantic knowledge (called “factual” for short): These were meaningful sentences bearing on generic semantic knowledge, i.e. facts of nature, culture, literature or the environment that are true or false independently of context. Some examples (translated into English) are “*The camel is a large humpbacked mammal” (True)*, “*The sun rises in the east every morning” (True), “Butterflies have small transparent wings” (False), or “Jupiter is the smallest planet in the solar system” (False).* The aim was to create a sharp contrast with syntactically anomalous sentences, thus activating brain regions involved in simple compositional semantics. In those sentences, generic world knowledge sufficed to decide whether the sentence was true or false. Of the 40 sentences, half of them were true (T), and half were false (F).
● Contextual knowledge: these sentences referred to facts in a specific context, for instance “*In Japan infant mortality is important”*. The context, which was always introduced by the first two or three words, was mostly geographical (a country, a city, a place, a planet…; 87.5 % of statements), but could also be historical (e.g. *in Greek mythology*…), literary (e.g. *in the Bible…*) or social (e.g. *During class*…). Intended as a control for social knowledge (see below), this condition forced participants to consider the specific context proposed, since the proposition’s truth value could not be immediately deduced from its out-of-context truth value (for instance, infant mortality may be low in general, but knowing this does not suffice to determine if it is low in Japan). Indeed, among the 20 sentences that were true, the main proposition was true out of context for half of them (sentences T/T), and false for the other half (T/F). And conversely, among the 20 sentences that were false, the main proposition was true out of context for half of them (F/T), and false for the other half (F/F). Therefore, there were 4 conditions with 10 statements each: true/true (T/T), true/false (T/F), false/true (F/T), and false/false (F/F). Some examples are: “*In Beijing pollution is a major problem” (T/T), “In volcanic islands the sand is black” (T/F), “In the Amazon spiders are harmless animals” (F/T),* and *“In Kenya the lakes of the plains freeze regularly” (F/F).* We hypothesized that adding semantic contextualization might activate additional semantic brain regions compared to mere factual knowledge (which it did).
● Social knowledge: These sentences referred to a person (or a group of people) and required that the person’s perspective and knowledge be considered in order to answer. As in the contextual knowledge condition, the truth value of the propositions often changed with the person’s point of view. Of the 40 sentences, half of them were overall true (T), and half were overall false (F). Among each, the principal proposition was true out of context for half of them (sentences T/T), and false for the other half (T/F). Thus, there were 4 conditions with 10 statements each: true/true (T/T), true/false (T/F), false/true (F/T), and false/false (F/F). Some examples are “*According to Darwin man is the cousin of the apes” (T/T), “For Mowgli the black panther is a friendly animal” (T/F), “For Jules Verne traveling to the center of the earth is impossible” (F/T)*, and *“For Einstein the atomic bomb is a harmless object” (F/F)*. The aim here was to maximize the differences with the contextual condition and elicit additional activation in the brain network consistently observed during Theory-of-Mind judgments (Deen et al., 2015a; Saxe & Kanwisher, 2003).
● Calculation: These sentences presented arithmetic calculations in verbal form, as a series of spoken or written words (no Arabic numerals were presented). Different numbers, ranging from 1 to 10,000 were allowed in this category, as long as they were expressed in four words or less. To increase the variability in this category, and match all stimuli for length, we also included words such as “double”, “half”, “one third” or “one quarter”. Some examples are “*Number three multiplied by twelve is thirty six” (T), “Half of two hundred is worth three hundred and fifty five” (F), or “The multiplication of four and three results in ten” (F).* Previous research has shown that calculation, compared to mere sentence processing, activates a distinct dorsal network, particularly in the intraparietal sulcus (e.g. Pinel et al., 2007); we aimed to replicate this result with semantic controls tightly matched for length and difficulty.
● Arithmetic principles: These sentences proposed abstract principles of arithmetic, defined as statements whose truth value does not depend on specific numerical values, and which therefore do not invite mental calculation, but only mathematical reflection on commutativity, neutral elements, parity, sign, etc. No numbers other than 0 and 1 were allowed in this category. Some examples are “*A positive number is larger than a negative number” (T)*, or *“A number multiplied by zero is equal to itself” (F)*. This condition was more exploratory since, to the best of our knowledge, arithmetic principles have not been previously dissociated from actual calculation; previous research has merely shown that, compared to non-mathematical statements, even abstract non-numerical mathematical statements yield activation in a broad math- and calculated-related network (e.g. Amalric & Dehaene, 2019).
● Geometry: These sentences stated geometric facts about basic shapes (squares, circles, rectangles, triangles, etc.) and features such as angles, parallelism, intersections, etc. Only the numbers one and two were allowed in this category. Some examples are “*A right angle is larger than an acute angle” (T)*, “*An equilateral triangle can be divided into two right triangles” (T)*, or *“A rectangle has only one axis of symmetry” (F)*. Our aim was to attempt to replicate previous suggestions of specific posterior intraparietal and inferior temporal contributions to geometry (Amalric & Dehaene, 2016).

All stimuli were created with the following general criteria: all verbs were in the present tense; there were no relative (subordinate) sentences; virtually all statements avoided negations (98.5% of sentences) and had only one verb (96.0%); or a maximum of 2 conjugated verbs). Non-mathematical stimuli did not contain any numbers (except for the word “*un”*, which is also an article in French) or other mathematical concepts such as measurements, geometric figures, etc. (although they did contain scientific statements such as “*Jupiter is the smallest planet in the solar system”(F)* or “*Dinosaurs are mammals who vanished from the earth”(F)*). All sentences were designed to be understood and answered correctly by a person with a level of education equivalent to the French National Diploma commonly called “brevet” (“*Diplôme National du Brevet des Collèges*”), which is passed at the end of 9th grade.

All stimuli were created in a written version (presented in rapid serial visual presentation [RSVP] on the screen) and in a spoken version (all recorded by a native speaker; all stimuli, including word lists, were recorded with a similar sentence-like prosody). As show in Figure S1, stimuli in the different categories were closely matched on the following criteria: number of words; total number of characters in written form (without blank spaces); mean word length (in letters); number of syllables (computed using this website: https://www.scribblab.com/outils/syllaber); mean log word frequency of occurrence in French (computed using http://www.lexique.org/); and duration of the spoken stimuli (measured using the “wave” library in python). Note that the duration of visual presentation was directly proportional to the number of words (350 ms/word). Figure S1 shows the corresponding means and standard errors.

### Task

The task was to indicate whether each statement was true or false. Participants responded by pressing a right-hand or left-hand button with their thumbs. For half of the participants, the right button press meant “true”, whereas the left button press meant “false”. For the other half of the participants, it was the opposite. Participants were instructed to answer as fast as possible after the end of each stimulus (similarly for the spoken and the written modalities). For stimuli without meaning (i.e. lists of words and meaningless sentences), they were instructed to respond “false”. All stimuli were randomly intermixed, and the participants were not told about the different categories before the experiment.

The experiment was programmed using Expyriment software (Krause & Lindemann, 2014)(http://expyriment.org). Response times were measured relative to the end of the auditory stimulus, or for visual stimuli, relative to the onset of the last word of the RSVP stream.

### Procedure

Before entering the scanner, the participants were familiarized with the task and were shown some examples of auditory and visual stimuli. 16 adult participants underwent five fMRI runs (full design), while 10 underwent only four runs. All 15 adolescents completed the full design. Each fMRI run lasted 9 minutes and 20 seconds, i.e. 64 stimulus epochs of 8 seconds, plus 8 randomly placed rest periods whose duration varied randomly between 4 and 8 seconds (mean = 6s in each run).

Each fMRI run comprised a pseudo-random mixture of visual and auditory stimuli of all types. The order of stimuli was randomized and was different for each subject, but there were always eight stimuli within each of the eight experimental conditions (lists of words, meaningless sentences, factual knowledge, contextual knowledge, social knowledge, calculations, arithmetic principles, geometry), for a total of 64 stimuli per run. Furthermore, in each category and subcategory, the number of auditory and visual stimuli, as well as of True (T) and False (F) sentences, was counterbalanced. Thus, for factual knowledge, calculations, arithmetic principles and geometry, there were four True and four False sentences; and for social and contextual knowledge, there were two T/T, two T/F, two F/T and two F/F sentences. Finally, for a given participant, within each run, half of the stimuli of each subcategory (T, F, T/T, T/F, F/T, F/F) were presented in written form and the other half in spoken form.

Written stimuli were presented using rapid serial visual presentation (RSVP, 350 ms per word; mean total duration = 3.2 s). Spoken stimuli had a mean duration of 3.13 s. Each stimulus was followed by a silent response period with a fixation cross, the duration of which was adjusted so that the stimulus onset asynchrony (SOA) was fixed at 8 s (figure 1). To facilitate the anticipation of stimuli, and since both auditory and visual stimuli were randomly intermixed, their onset and their modality were announced, 350 ms before stimulus onset, by a red cross for written stimuli and by a beep for spoken stimuli.

**Figure 1.**
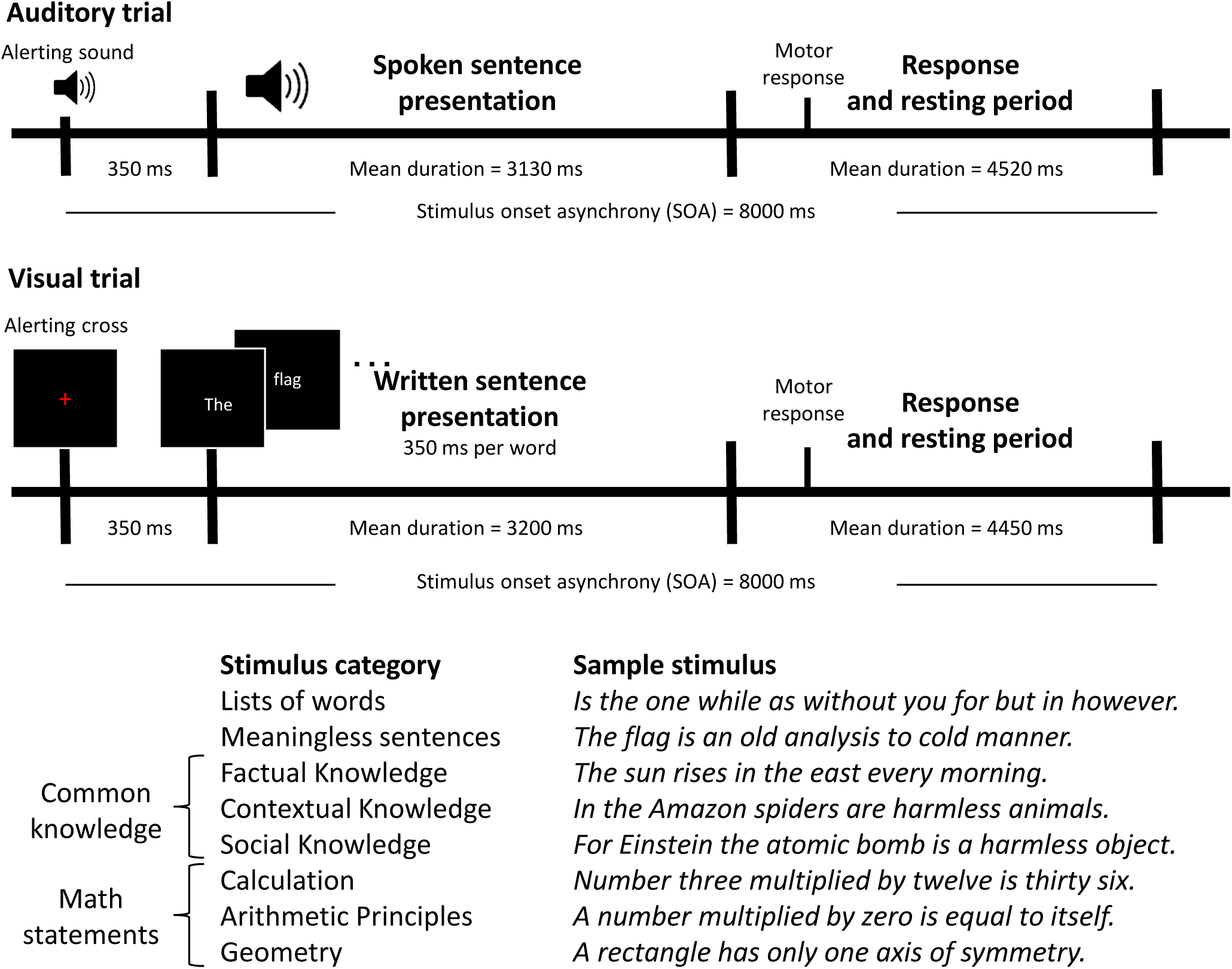
Experimental design and examples of stimuli in each category. In this event-related 3T fMRI paradigm, participants were asked to judge short statements as true or false. They heard or read a mixture of auditory and visual statements ranging from lists of words to meaningless sentences (to which they were asked to always respond ‘false’) and various meaningful math and non-math statements, half of which were true.

### MRI data acquisition

MRI data was acquired on a 3 Tesla Siemens Prisma system equipped with a sixty-four channel coil. For each participant, one anatomical image was acquired using a 3D gradient-echo sequence and a voxel size of 1.0×1.0×1.0 mm^3^ (for 16 of the adult participants and all 15 adolescents) and of 0.9 x 0.9 x 0.9 mm^3^ (for the other adults (n=10)). Functional images were acquired during 5 sessions (for 16 adult participants and 15 adolescent participants) or 4 sessions (for 10 adults) with a number of scans ranging from 275 to 288 each, using a sequence with the following parameters. For the main group of 16 adult participants and for the adolescents, we used the Minneapolis BOLD cmrr_mbep2d sequences (Moeller et al., 2010; Setsompop et al., 2012), with TR = 2.1 s TE = 30.4 ms, matrix = 110×110, voxel size = 1.75×1.75×1.75 mm^3^, 78 slices in interleaved order, multiband acceleration factor = 3, iPAT0, partial Fourier = 7/8, number of scans = 275. The 10 other adults were scanned with Minneapolis BOLD cmrr_ep2d sequences (Moeller et al., 2010; Setsompop et al., 2012), with TR = 2 s, TE = 26.80 ms, matrix = 128×128, voxel size = 1.5×1.5×1.5 mm^3^, 93 slices in interleaved order, multiband acceleration factor = 3, iPAT2, partial Fourier deactivated, number of scans = 280 to 288.

### Data processing and analysis

Data processing was performed using SPM12 (http://www.fil.ion.ucl.ac.uk/spm/, Welcome Department of Cognitive Neurology). The anatomical scan was spatially normalized to the Montreal Neurological Institute avg152 T1-weighted brain template using default parameters (including nonlinear transformations and trilinear interpolation). Functional volumes were corrected for slice timing differences (first slice as reference), realigned after motion correction (registered to the mean using 2^nd^ degree B-Splines), corrected for distortion (using the “topup” algorithm (Andersson et al., 2003) as implemented in FSL (Smith et al., 2004), coregistered to the anatomy (using Normalized Mutual Information), spatially normalized using the parameters obtained from the normalization of the anatomy, and smoothed with an isotropic Gaussian kernel (FWHM=5mm).

Experimental effects at each voxel were estimated using a multi-session design matrix modeling, for each run, the 18 conditions (8 categories x 2 modalities + separate regressors for the right and left motor responses), plus the 6 motion parameters computed in the realignment phase. To permit comparison across conditions, stimulus-evoked responses were modeled by a fixed hemodynamic response function (HRF), obtained by convolving a 3.2 s rectangular kernel with the standard SPM impulse HRF, as well as its time derivative. The two regressors for motor responses were obtained by convolving the same HRF with an arbitrarily short (100 ms) rectangular impulse.

Contrasts for each condition relative to baseline were computed by averaging the regression coefficients across fMRI runs. Individual contrasts were smoothed with an 8×8×8mm Gaussian kernel before entering them into a second-level one-way analysis of variance model with one regressor per experimental condition and one regressor per subject. Comparisons between conditions were then performed at this second level, with a voxelwise statistical threshold of p<0.001, and a clusterwise size threshold of p<0.05 FWE corrected for multiple comparisons over the entire brain volume.

The results shown in the figures were created using the bspmview toolbox (DOI:10.5281/zenodo.168074).

## Results

### Behavioral results

All adults (n=26) performed the task with relatively low error rates (mean ER = 12.8%; standard error [SE] = 1.30% standard error, range 4.69%-29.8%) and a response time of 1.21s (SE = 0.06s) following sentence ending. ANOVAs indicated that sentence category (8-level factor), presentation modality (2-level factor: written/spoken), and their interaction influenced both error rates (category: F(7,175) = 36.4, p < 10^-15^; modality: F(1,25) = 9.43, p = 0.005; interaction: F(7,175) = 3.23, p = 0.003) and response time (category: F(7,175) = 33.9, p < 10^-15^; modality: F(1,25) = 24.7, p < 10^-4^; interaction: F(7,175) = 5.24, p < 10^-4^).

As can be seen in figure 2, unsurprisingly, the default ‘false’ responses to word lists and meaningless sentences were faster and more accurate than the more thought-demanding responses to meaningful statements (t(25) = 6.77 and 12.27, respectively, p < 10^-6^). Within the six meaningful conditions, there was no main difference in error rate between mathematical and factual knowledge (t(25) = 0.97, p = 0.339), but mathematical statements elicited a longer response time than factual knowledge (t(25) = 5.44, p < 10^-4^). However, this was not true within each condition, since the calculation condition was systematically less error-prone than the non-math conditions (t(25) = 4.24, p < 0.0003) and did not differ in response time. In descending order of response time, the slowest category was *Geometry* (RT = 1.53 ± 0.069 s), followed by *Arithmetic Principles* (RT = 1.40 ± 0.072 s), *Social Knowledge* (RT = 1.19 ± 0.080 s), *Contextual Knowledge* (RT = 1.19 ± 0.065 s), *Factual Knowledge* (RT = 1.18 ± 0.059 s), and *Calculation* (RT = 1.17 ± 0.069 s). Thus, activations that would be systematically higher for every category of mathematical statements cannot be attributed to a difficulty effect.

**Figure 2.**
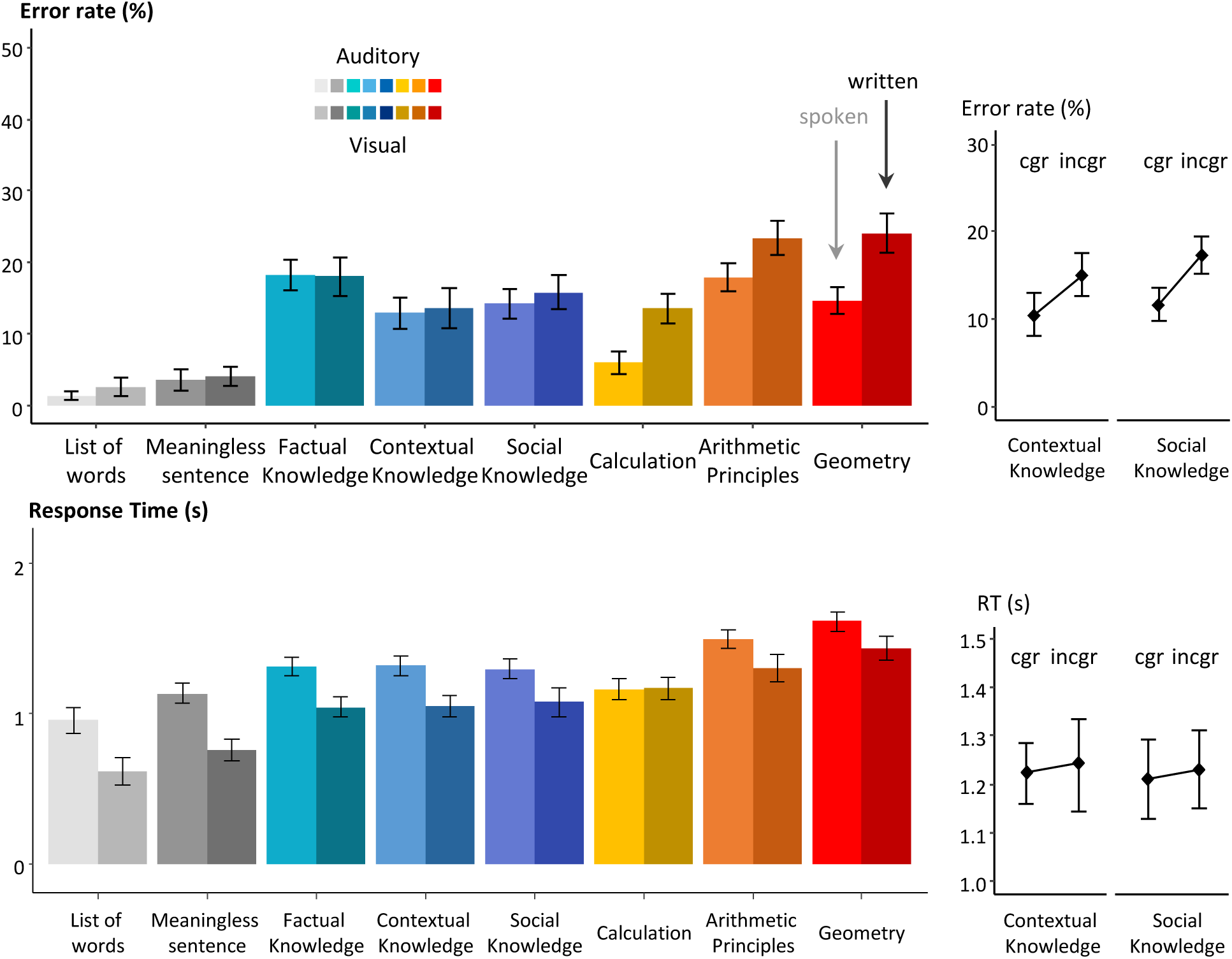
Behavioral results for adults. Average error rates (top) and response times (RT, bottom) revealed that (a) lists of words and meaningless sentences were responded quickly and accurately with a default “false” response; (b) meaningful statements varied in difficulty, yet without any systematic difference between math and non-math statements; (c) RTs (measured relative to sentence ending) were slower on auditory than on visual trials; (d) For contextual and social statements, incongruent trials (in which the truth value of the full statement differed from that of its main proposition, e.g. “*For Donald Trump, climate change is worrying*”) led to more errors than congruent trials (e.g. *“In Beijing, pollution is a major problem”)*.

An ANOVA restricted to meaningful statements, with category, modality, and truth value as factors, found main effects of modality on both error rate (F(1,25) = 9.89, p = 0.004) and response time (F(1,25) = 18.1, p < 0.001), indicating a speed-accuracy trade-off: responses to written statements were faster than to spoken statements, but more error-prone. More interestingly, modality interacted with math/non-math status (errors: F(1,25) = 9.79, p = 0.004; RTs: F(1,25) = 8.83, p = 0.006). Written mathematics induced more errors than spoken mathematics (F(1,25) = 20.9, p < 0.001), while there was no such modality effect on non-mathematical statements (F(1,25) = 0.164, p = 0.689). For RTs, only the calculation condition differed from the others in that it did not show faster responses for written than for spoken statements. Overall, mathematical statements seemed to suffer more than non-mathematical statements from a presentation in the visual modality – a pattern that, as we shall further discuss later, is compatible with an interference of visual sentences on the visual areas recruited by mathematics.

The truth value of the statements affected both error rates (F(1,25) = 12.1, p = 0.002), and response times (F(1,25) = 24.3, p < 10-4), indicating that false statements were more difficult to judge than true ones. Truth value interacted significantly with category (ER: F(5,125) = 10.5, p < 10-7; RTs: F(5,125) = 12.4, p < 10-9), mainly due to an increased difficulty for false *arithmetic principles* and *geometry* statements. Finally, truth value did not interact with presentation modality on error rates (F(1,25) = 1.20, p = 0.284), but did on response times (F(1,25) = 5.27, p = 0.030), due to a reduced effect of modality on response times for false statements compared to true statements.

For contextualized sentences only (i.e. *Contextual Knowledge* and *Social Knowledge*), we evaluated the effect of congruity between the core statement and the context. Remember that half of the stimuli were congruent statement, where the truth value of the sentence was unchanged by the particular geographic, temporal or personal context; while the other half were incongruent statements whose true value flipped in the particular context. We conducted 4-way ANOVAs with factors of congruity, truth value, modality, and category (Contextual vs Social). While no effect was found for RTs, error rates were affected by a main effect of congruity (F(1,25) = 9.80, p = 0.0043): participants made more errors when the context induced a change in truth value (figure 2, right column). This finding confirms our hypothesis that the contextual and social conditions, while not overall more difficult than the others, involved an additional computation to overcome general sentential meaning and focus on the specific context. There were no interactions of congruity with any of the other factors, suggesting that this semantic effect occurred at a modality-independent level and that the Social and Contextual conditions were well matched in this respect.

For the sake of brevity, we do not report the adolescent data in the same level of detail here. As shown in figure 3, the results were virtually identical. In particular, we replicated (1) the fact that mathematical statements were not systematically more difficult than non-math statements (as in adults, calculation was the easiest), and (2) a modality X category interaction (F(5,60) = 4.02, p = 0.003 on RTs) suggesting that math suffers more than non-mathematics from being presented in the visual rather than the auditory modality. The only potentially significant difference with adults was a small unexplained congruity X category interaction: adolescents, unlike adults, showed a congruity effect on errors only in the social but not in the contextual condition.

**Figure 3.**
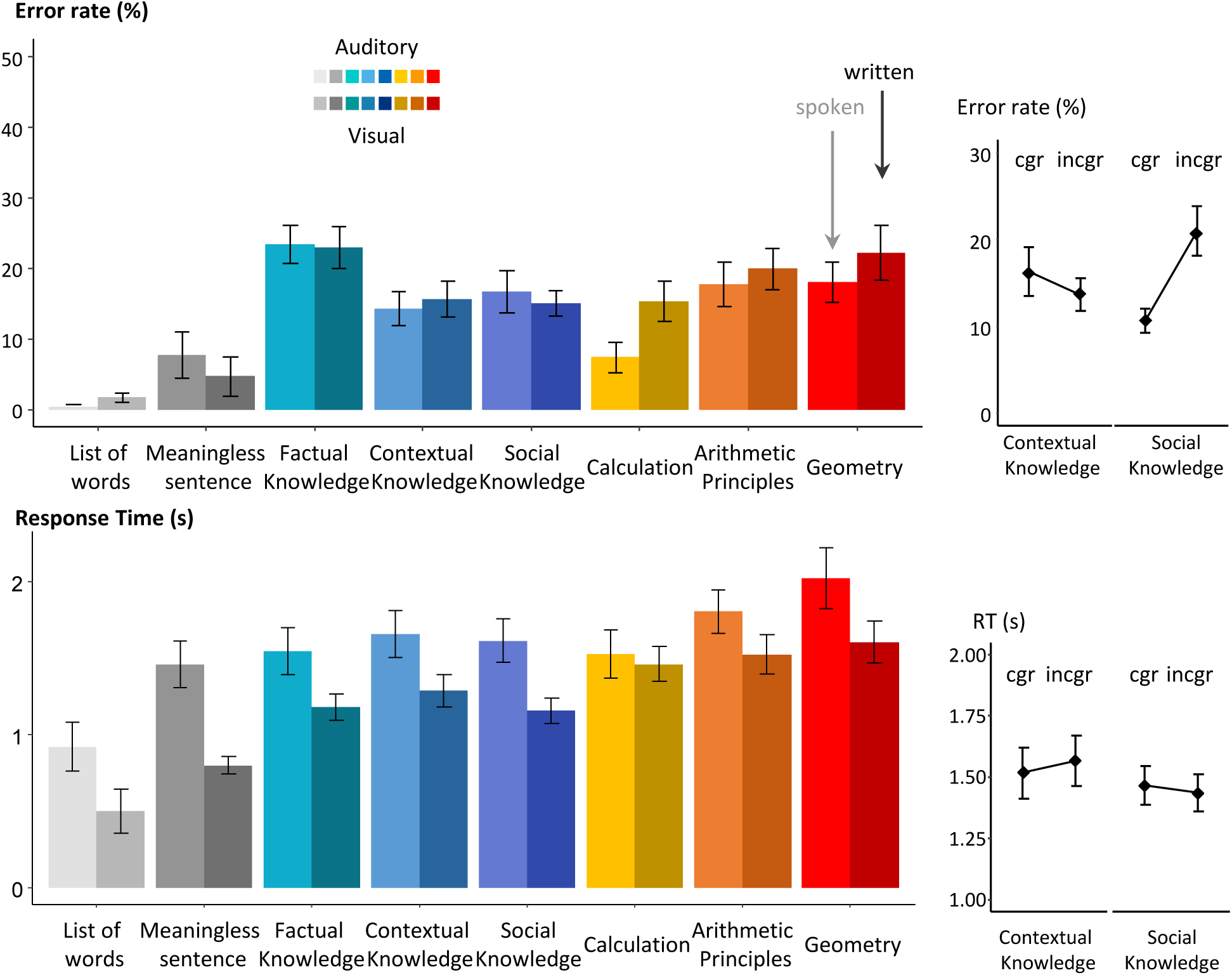
Behavioral results for adolescents. Average error rates (top) and response times (RT, bottom) revealed that (a) lists of words and meaningless sentences were responded quickly and accurately with a default “false” response; (b) meaningful statements varied in difficulty, yet without any systematic difference between math and non-math statements; (c) RTs (measured relative to sentence ending) were slower on auditory than on visual trials; (d) For social statements, incongruent trials (in which the truth value of the full statement differed from that of its main proposition, e.g. “*For Donald Trump, climate change is worrying*”) led to more errors than congruent trials. However, unlike for adults, for contextual statements, incongruent trials led to fewer errors than congruent trials.

### FMRI results

#### Brain networks for syntactic structure in meaningless sentences

The first level of our hierarchical design contrasted meaningless sentences and lists of words, presumably isolating the stage of syntactic composition while reducing the possibility of semantic composition. This contrast, shown in figure 4A, isolated a left-lateralized network of areas involving (1) much of the left superior temporal sulcus (STS), including both anterior and posterior sites, (2) the left inferior frontal gyrus (IFG), including both pars triangularis and orbitalis (but not opercularis). Smaller activations were seen in the right IFG and right cerebellar Crus II (CerCrus2).

**Figure 4.**
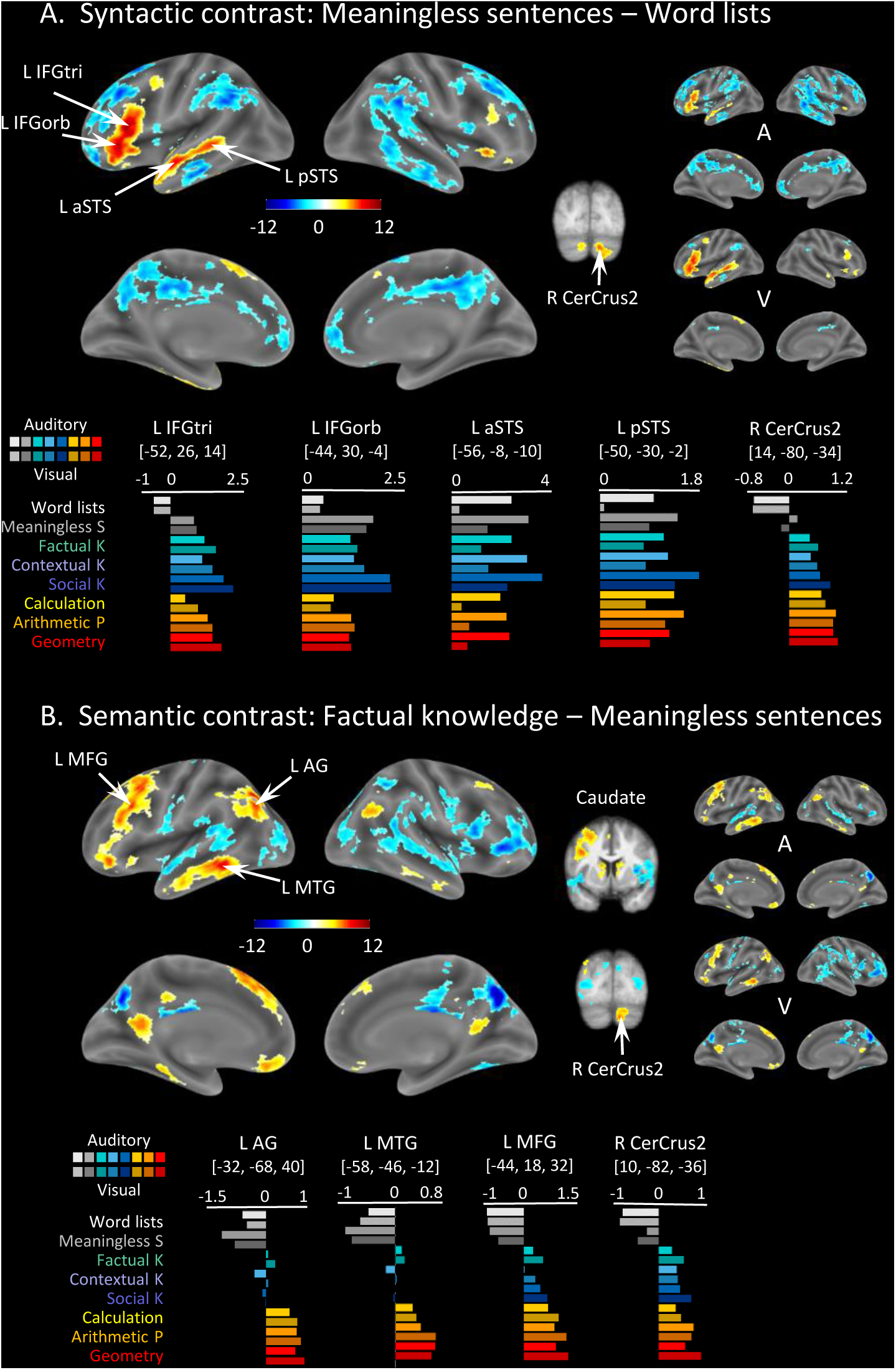
fMRI parsing of the language network. A, Syntactic contrast: greater activity for meaningless sentences than for word lists (red) or the converse (blue). B, Semantic contrast: greater activity for meaningful sentences expressing factual knowledge than for meaningless sentences. All images are thresholded at voxelwise p<0.001 and FWE-corrected cluster-level p<0.05). Inflated-brain images show the main contrast pooling over both modalities (left), and the separate contrasts for auditory (A) and visual (V) modalities (smaller brains at right). Histograms show the mean fMRI activation (beta value) at selected peaks for each of the 16 stimulus categories. Abbreviations: L = Left, R = Right, IFGtri = Inferior Frontal Gyrus *pars triangularis*, IFGorb = Inferior Frontal Gyrus *pars orbitalis*, aSTS = anterior Superior Temporal Sulcus, pSTS = posterior Superior Temporal Sulcus, CerCrus2 = Cerebellum Crus II, MFG = Middle Frontal Gyrus, AG = Angular Gyrus, MTG = Middle Temporal Gyrus, A = Auditory, V = Visual.

Figure 4A shows histograms of activations for all conditions. These plots clearly show that in these regions, meaningless sentences as well as all the other sentences led to strong activations, while only the word list condition led to reduced activation or even to a deactivation relative to rest (in L IFGtri and R CerCrus2). All these regions showed a significant difference separately for auditory and visual stimuli (Fig. 4A, right insets A for auditory, V for visual) and showed no significant interaction with modality. Indeed, in IFG and CerCrus2, the activation profiles were strictly independent of modality, confirming their contribution to amodal processes of phrase integration. In STS, there was an additional main effect of modality, with an overall greater activation for auditory than for visual sentences. This effect could be due to a partial volume effect related to the proximity of auditory areas in the STG, a possibility that could be addressed by high-resolution single-subject imaging.

A significant interaction with modality (V>A) was found only in a few distinct regions: the superior temporal gyrus (STG), especially in the right hemisphere, as well as the right IFG and the middle frontal gyrus (MFG), suggesting that the visual modality of presentation led to a larger difference between meaningless sentences and word lists in these areas (Figure S2A), perhaps due to greater difficulty due to the fast pace of visual presentation.

### Brain networks for factual semantic knowledge

The second level of our hierarchy contrasted meaningful sentences, which refer to simple factual semantic knowledge with meaningless sentences, presumably isolating processes of elementary semantic composition and truth-value judgment. This contrast isolated a new set of brain regions, again predominantly lateralized to the left hemisphere (Figure 4B): the left angular gyrus (AG), a large part of the left middle temporal gyrus (MTG) and of the left middle frontal gyrus (MFG), extending into the mesial prefrontal cortex (mPFC) as well as the orbitofrontal cortex (OFC), the left retrosplenial cortex (RSC), and the right CerCrus2. As shown in figure 4B, histograms showed a very low level of activation or even deactivation in these regions for both word lists and meaningless sentences, and a sudden jump to a higher level when meaningful sentences appeared (further differences between conditions are discussed below).

All these regions were activated identically in the visual and auditory modalities, and none showed a significant interaction with modality (the only significant interaction occurred in a small bilateral superior frontal region with greater activation in the auditory than in the visual modality; MNI coordinates: 18, 22, 50 and −18, 12, 56, see figure S2B).

### Brain networks for contextual and social knowledge

We then examined which brain areas, if any, were further modulated by the need to contextualize semantic knowledge to a specific situation (e.g. “In Japan,…”), and whether some areas were specific to the social context (e.g. “According to Einstein,…”). These contrasts are shown in figure 5.

**Figure 5.**
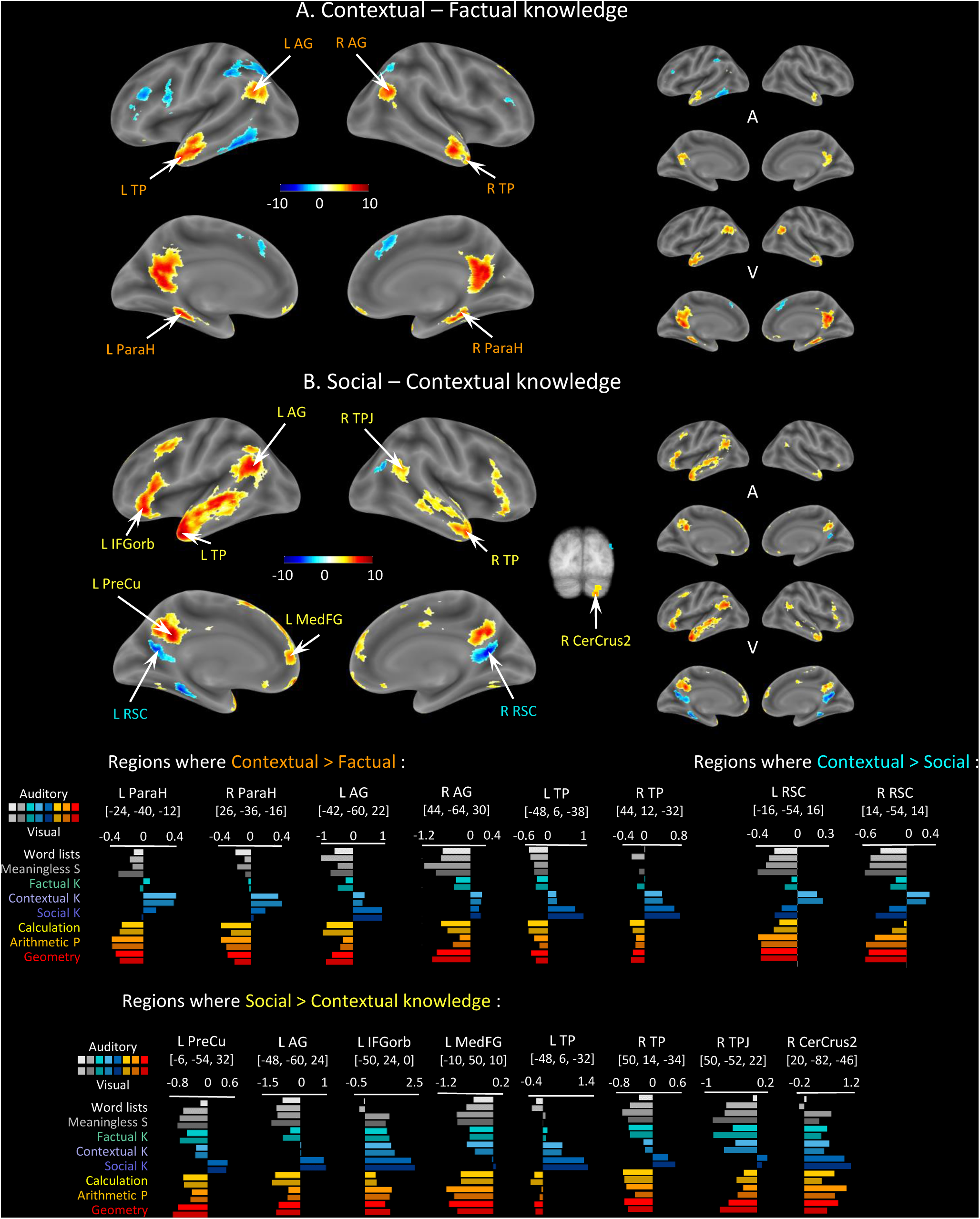
Brain areas involved in contextual and social statements. Same format as figure 4. A large extent of the classical fronto-temporal language network shows greater activation to social knowledge statements (“According to Einstein, etc”) than to matched contextual statements of similar length and syntax, yet bearing on geographic rather than human context (“In volcanic islands, etc.”) (panel B, and last row of histograms). Many of these regions also show greater activation to contextual statements relative to plain factual ones (panel A), but some regions show specificity for social knowledge (e.g. right TPJ, precuneus) and others for contextual statements (bilateral retrosplenial cortex and parahippocampal cortex). Abbreviations: ParaH = Parahippocampal Gyrus, TP = Temporal Pole, TPJ = Temporo-Parietal Junction, MedFG = Medial Frontal Gyrus, RSC = Retrosplenial Cortex, PreCu = Precuneus.

Sentences requiring contextual knowledge, relative to factual knowledge, elicited bilateral activation in temporal poles (TP), angular gyri (AG), retrosplenial cortex/precuneus (RSC/PreCu), and parahippocampal gyri (ParaH) (Figure 5A). These activations were symmetrical in the two hemispheres. Histograms clearly showed how these regions remained silent or even deactivated in most conditions (including meaningful but mathematical materials), but suddenly and selectively activated to sentences that required a contextualization of knowledge. None of these regions showed any influence of the visual or auditory modality.

Compared to this contextual condition, did social sentences evoke any additional brain activity? The answer was positive in a large set of brain regions with a clear left hemisphere lateralization (Figure 5B). These regions could be divided into two groups: first, some areas were already activated by meaningless sentences (in L STS, L IFGorb), factual or contextual knowledge (in L AG, bilateral TP and PreCu, R CerCrus2), but showed even more activity in response to social sentences; second, to a much lesser extent, some areas seemed to be specifically activated by social sentences. This effect was primarily seen in the right temporo-parietal junction (TPJ) (Saxe & Kanwisher, 2003), but also at sites in the mesial frontal cortex (medFG), left AG and precuneus. As shown in the histograms at the bottom of figure 5, the R TPJ and L PreCu, were deactivated by all conditions except for the social sentences. Again, these regions were modality-independent.

### Math-responsive areas

Compared to the three meaningful non-mathematical conditions (factual, contextual, and social), the three mathematical conditions induced substantial differences in brain activity (figure 6). On the positive side, mathematical sentences induced greater activity in a large bilateral network that has previously been described as math-responsive (Amalric & Dehaene, 2016, 2017, 2019) and includes sites in bilateral intraparietal sulci (IPS), inferior temporal gyri (ITG), middle frontal gyri (MFG), extending into the dorsal part of the IFG opercularis (area 44d), and posterior cingulate (PostCing), as well as the caudate and the Culmen region of the cerebellum.

**Figure 6.**
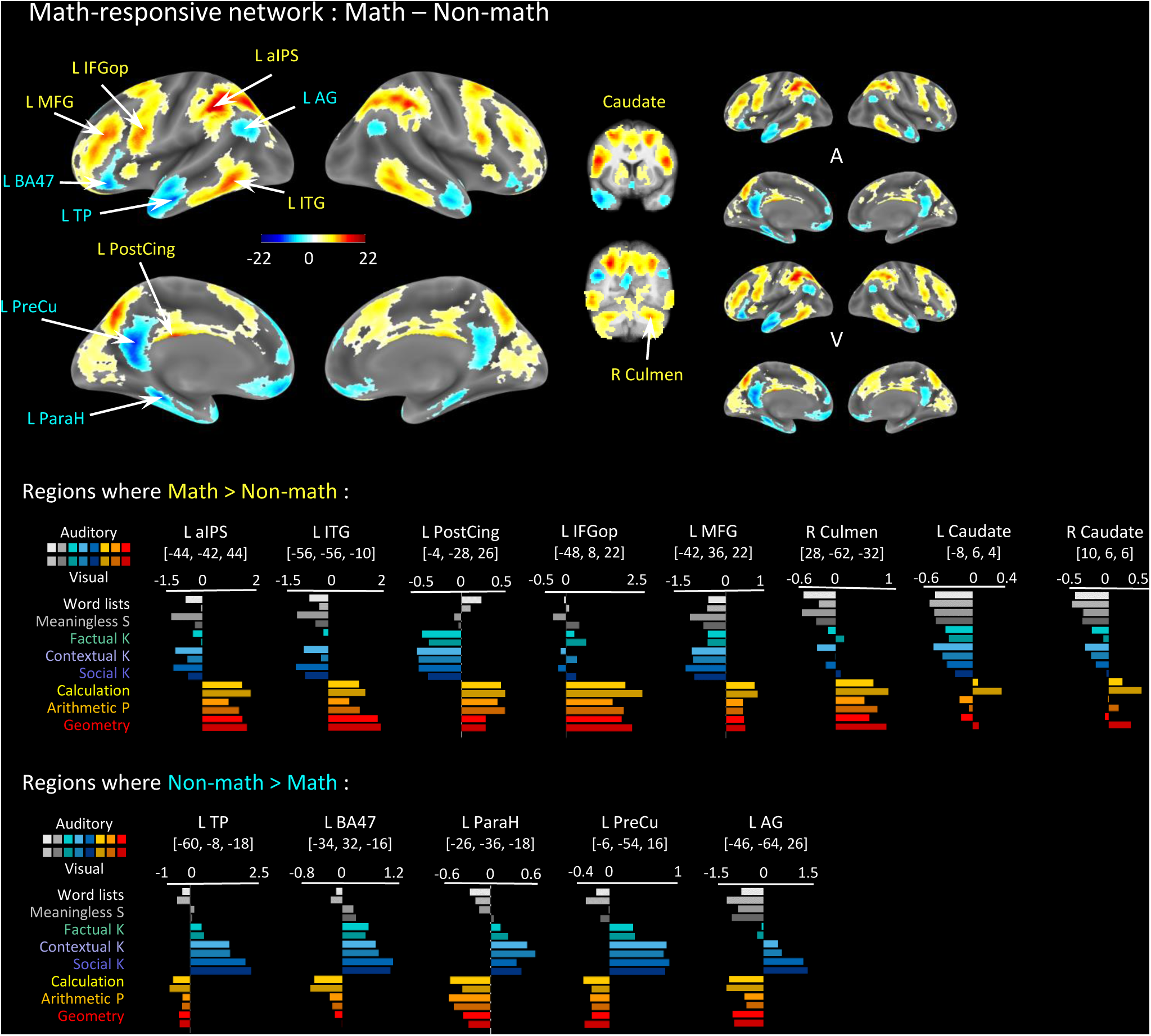
Math-responsive network, identified by the contrast of all math statements relative to all non-math statements. Same format as figure 4. Regions in red show greater activity for math than for control non-math statements; their activation profiles at the main peaks are plotted in the first line of histograms below. Regions in blue show greater activity for meaningful non-math statements and are plotted in the second line of histograms. Abbreviations: IFGop = Inferior Frontal Gyrus *pars opercularis*, aIPS = anterior Intraparietal Sulcus, BA47 = Brodmann Area 47, ITG = Inferior Temporal Gyrus, PostCing = Posterior Cingulate Cortex.

Conversely, mathematics led to reduced activity in most of the previously described bilateral semantic network, including bilateral AG, TP, inferior frontal BA47, precuneus/retrosplenial, and parahippocampal areas. As shown in the histograms of figure 6, these regions remained silent (at the same level as word lists) or even deactivated by mathematical sentences. Conversely, the math-responsive regions that were strongly activated by all mathematical statements, in both visual and auditory modalities, were not activated or even deactivated by non-mathematical sentences. Thus, a double dissociation between math and non-math was observed, extending previous findings (Amalric & Dehaene, 2016, 2017, 2019).

There were no interactions with modality across the entire math-responsive network. The only exceptions were a small right mid/anterior cingulate site with greater activation for math in the visual modality (MNI: 6, 30, 34; see figure S2C), and conversely a small cerebellar site of greater activation for math in the auditory modality (MNI: 18, −74, −38, corresponding to Crus2; see figure 5 bottom right for a histogram). These might be false positives, as they were not replicated in the adolescent group (figure S3C).

We next searched for differences among the three types of math statements (see figure 7). Calculation was the most distinct condition, with greater activation relative to the other two math conditions in bilateral horizontal IPS (hIPS), precentral/IFGop, SMA, anterior insula, and bilateral spots in the anterior putamen. Arithmetic principles yielded relatively more activity in left MFG and left AG (with smaller effects in right MFG and AG), left MTG and left BA11. Curiously, while largely silent for non-math sentences, these regions also activated in response to social knowledge, perhaps suggesting a role in multiple forms of abstract high-level reasoning (figure 7, left bottom row).

**Figure 7.**
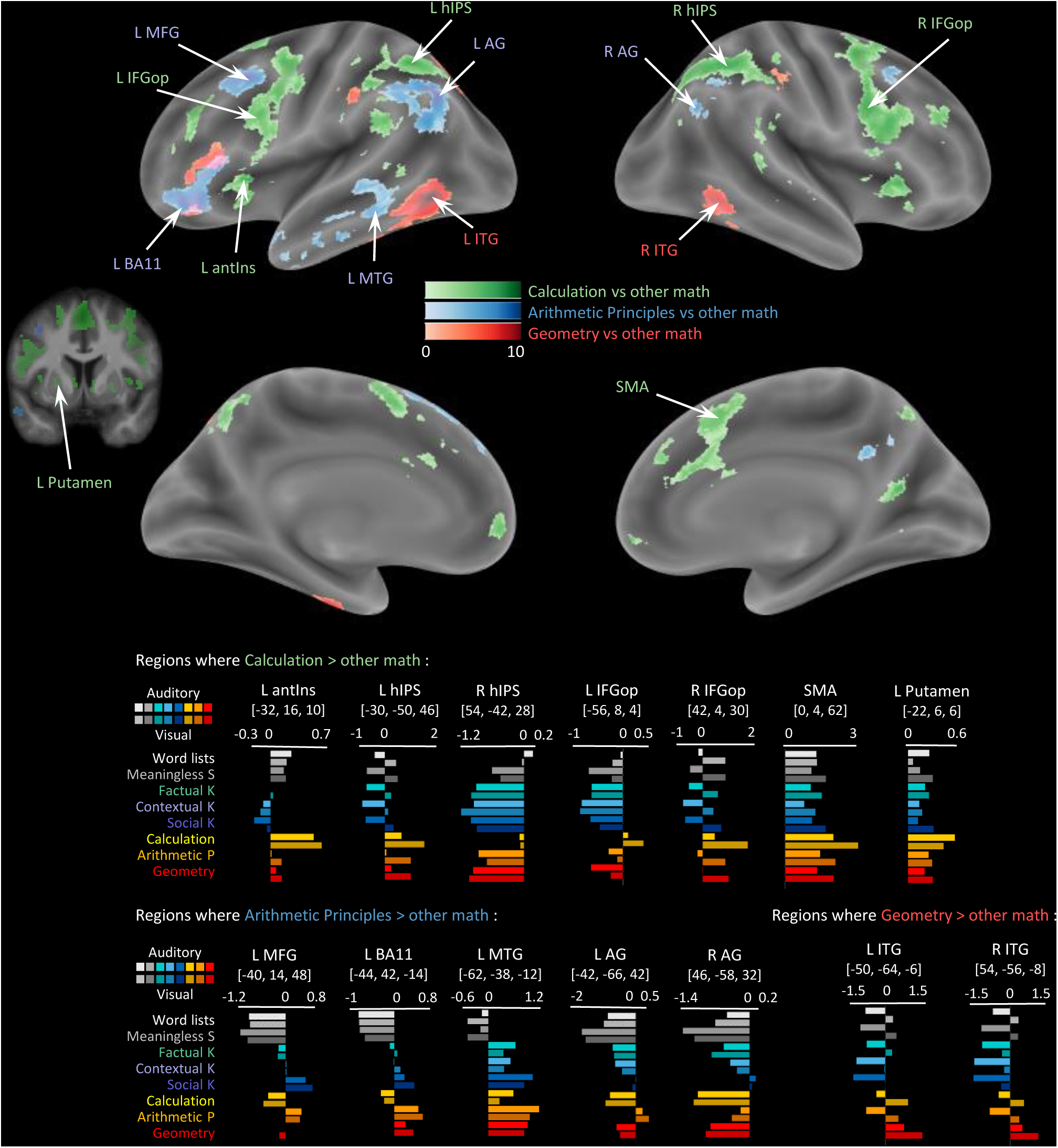
Differences between the three domains of mathematics. Each color shows the result of a contrast for one type of math statement (e.g. calculation statements) versus the average of the other two (here, arithmetic principles and geometry). To focus only on positive activations, images are masked by the contrast of the appropriate math condition (e.g. calculation) relative to wordlists. All images are thresholded at voxelwise p<0.001 and FWE-corrected cluster-level p<0.05). The main results are (1) greater activation for calculation (green) in a large extent of the overall math-responsive network (bilateral IPS, IFG, anterior insula, SMA); (2) greater activation for arithmetic principles (blue) in a left-lateralized network involving angular gyrus, MFG and ventral frontal cortex (BA11); (3) greater activation for geometry (red) primarily in bilateral inferior temporal cortex. Abbreviations: hIPS = horizontal segment of the Intraparietal Sulcus ((Dehaene et al., 2003)), BA11 = Brodmann Area 11, antIns = anterior Insula, SMA = Supplementary Motor Area.

Finally, geometry yielded greater activation in the left and right posterior sectors of the ITG. Interestingly, those activations were located at the posterior tip of the large ITG region activated by all mathematical statements (figure 6). Furthermore, they also showed a main effect of modality (greater activity by visual than by auditory statements). Thus, the geometrical content of sentences, perhaps because of its greater visual imagery content, activated a posterior math-responsive inferior temporal region just anterior to occipital cortex.

When analyzing interactions with the modality of sentence presentation, the same bilateral posterior ITG region showed up in two contrasts. First, the contrast “geometry > other math” in bilateral fusiform gyrus/pITG was stronger when sentences were presented auditorily than visually (peak MNI coordinates: −40, −58, −6 and 48, −70, 0; see figure S2D). Second, conversely, calculation, relative to other math, evoked more activity in the visual than in the auditory modality, in bilateral inferior temporal/occipital gyri (44, −72, 2; −44, −74, −4) and fusiform gyri (44, −48, −20; −38, −54, −16; see figure S2E). This complex pattern suggests that the modality of stimulus presentation may have interfered with the contents of the sentences, in agreement with our behavioral findings that visual presentation slowed down the resolution of calculation problems while accelerating the resolution of geometry problems (see Figure 2). These findings, however, should not detract from the main observation that most of the math-responsive network, including a slightly more anterior region of pITG responded strongly to all math statements in both visual and auditory modalities (figure 6).

### Replication in adolescents

As mentioned in the introduction, one of the goals of our study was to develop a stable, highly replicable paradigm for evaluating the integrity of language, math and social knowledge brain networks in diverse populations, including individuals with developmental conditions. To evaluate the replicability of our results, we scanned 15 adolescents with the same exact stimuli and conditions as in the adult group. The corresponding results are presented in figures 8-11, following the same format as figures 3-7. Figure 12 shows the results of an ANOVA comparing the fMRI results of the 26 adults and 15 adolescents. Despite the smaller sample size in the adolescent group, all major findings were successfully replicated. Briefly:

● Meaningless sentences, relative to word lists, activated a core language network primarily comprising left STS and IFG (figure 8A).
● Factual sentences yielded an additional activation in AG, MTG, and CerCrus2. Although most of the MFG remained below threshold in this contrast in adolescents (Figure 8B).
● Histograms at the adult peak showed that it was, in fact, activated, without any significant difference between groups. The only significant interaction was a slightly increased activation in mesial superior frontal gyrus in adults (figure 12A).
● Contextual versus factual knowledge activated the same bilateral areas as in adults, including the AG, TP, and parahippocampal cortex (figure 9A).
● Social, relative to contextual knowledge, again activated a large chunk of the left-hemisphere STS and IFGorb, precuneus, and small foci in right TPJ and TP (figure 9B). These regions did not differ significantly between the two groups, except for a slightly larger activation in a right anterior temporal/TP cluster in adults (figure 12B).
● Mathematics, relative to non-math statements, activated the same set of bilateral regions in IPS, ITG, MFG, IFGop, posterior cingulate and Culmen (figure 10). While the overall math-responsive network was largely identical across adults and adolescents, significant group-by-condition interactions were observed. Specifically, several regions in left temporal lobe, left IFG and right TP, were more activated in adults; and conversely, posterior cingulate and midline frontal regions were more activated in adolescents (figure 12C). Notably, these interaction effects did not overlap with the core math-responsive network and appeared to stem primarily from greater math-related deactivation in adolescents, complicating their interpretation. Finally, the comparisons of the three math conditions largely replicated the adult results, with calculation yielding more activity in bilateral IPS, IFGop and anterior insula; arithmetic principles in left AG and TP; and geometry in bilateral posterior ITG. No group interactions were found on these contrasts (figure11).

**Figure 8.**
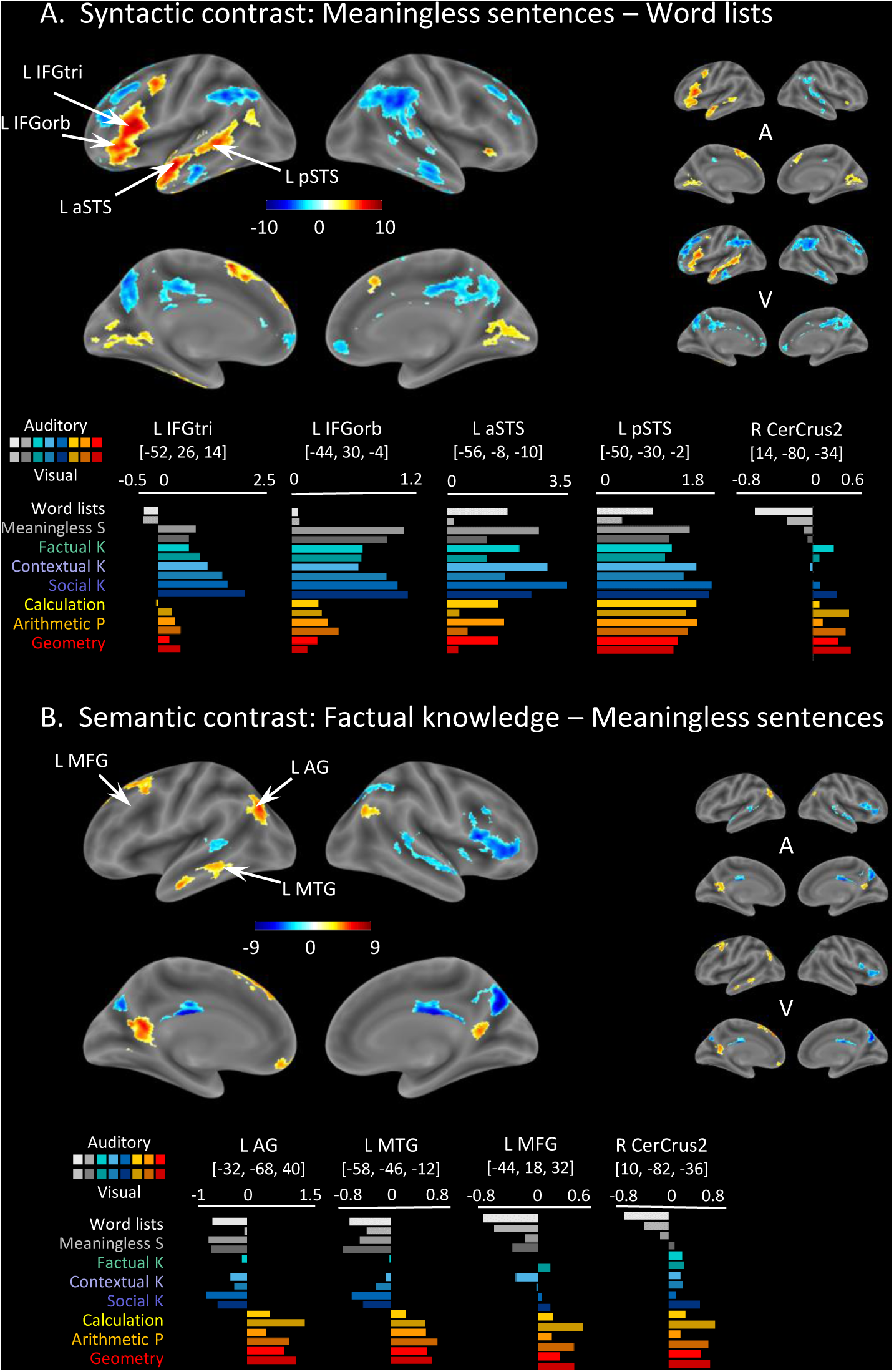
fMRI parsing of the language network in adolescents. Same format as figure 4. A, Syntactic contrast: greater activity for meaningless sentences than for word lists (red) or the converse (blue). B, Semantic contrast: greater activity for meaningful sentences expressing factual knowledge than for meaningless sentences. All images are thresholded at voxelwise p<0.001 and FWE-corrected cluster-level p<0.05). Inflated-brain images show the main contrast pooling over both modalities (left), and the separate contrasts for auditory (A) and visual (V) modalities (smaller brains at right). Histograms show the mean fMRI activation (beta value) at selected peaks for each of the 16 stimulus categories.

**Figure 9.**
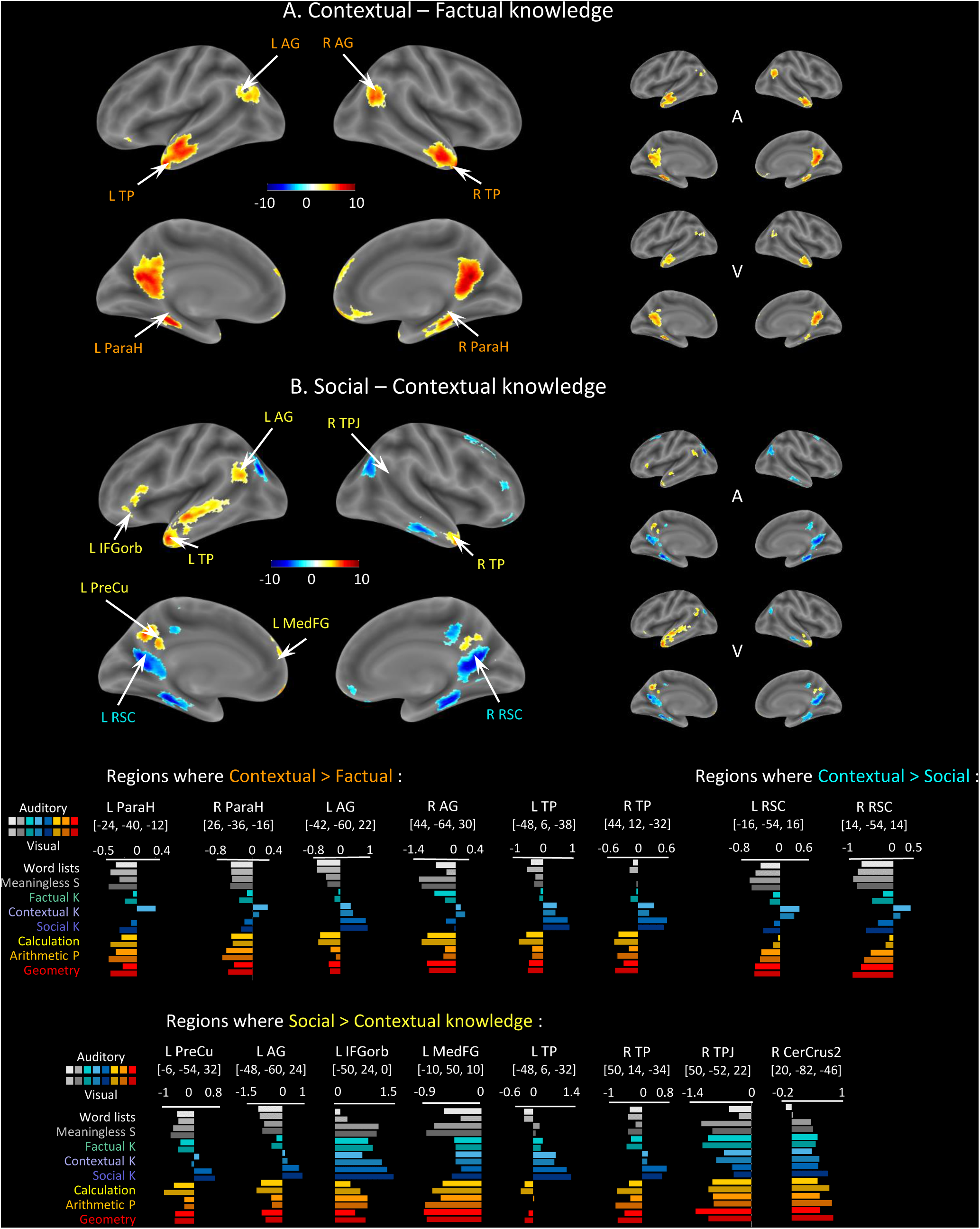
Brain areas involved in contextual and social statements in adolescents. Same format as figure 4. A large extent of the classical fronto-temporal language network shows greater activation to social knowledge statements (“According to Einstein, etc”) than to matched contextual statements of similar length and syntax, yet bearing on geographic rather than human context (“In volcanic islands, etc.”) (panel B, and last row of histograms). Many of these regions also show greater activation to contextual statements relative to plain factual ones (panel A), but some regions show specificity for social knowledge (e.g. precuneus) and others for contextual statements (bilateral retrosplenial cortex and parahippocampal cortex). The social network appears to be less extended for adolescents than for adults (e.g. right TPJ does not appear here but see figure 12 for direct comparisons between adults and adolescents).

**Figure 10.**
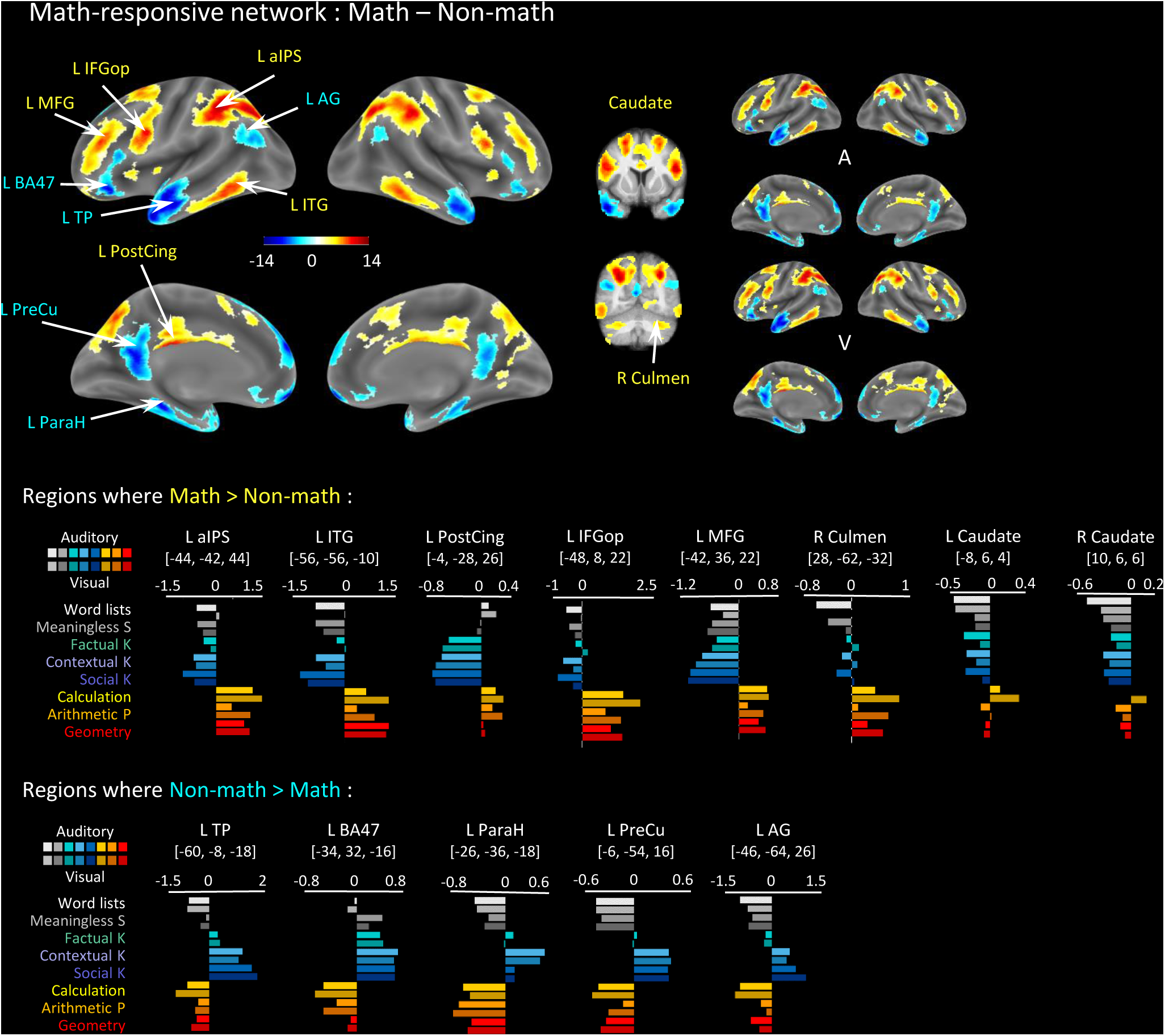
Math-responsive network in adolescents, identified by the contrast of all math statements relative to all non-math statements. Same format as figure 5. Regions in red show greater activity for math than for control non-math statements; their activation profiles at the main peaks are plotted in the first line of histograms below. Regions in blue show greater activity for meaningful non-math statements and are plotted in the second line of histograms.

**Figure 11.**
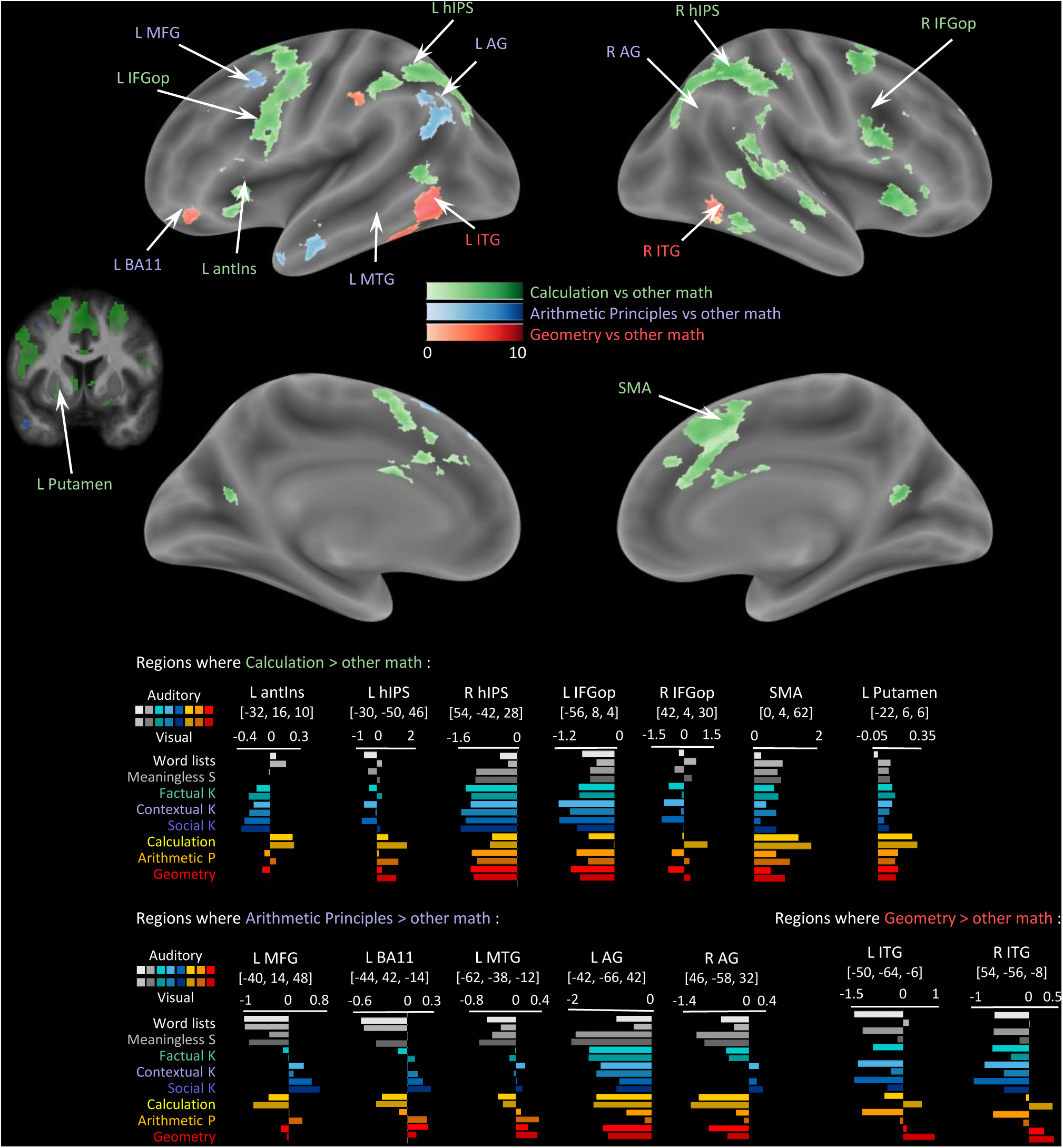
Differences between the three domains of mathematics in adolescents. Same format as figure 6. Each color shows the result of a contrast for one type of math statement (e.g. calculation statements) versus the average of the other two (here, arithmetic principles and geometry). To focus only on positive activations, images are masked by the contrast of the appropriate math condition (e.g. calculation) relative to wordlists. All images are thresholded at voxelwise p<0.001 and FWE-corrected cluster-level p<0.05). The main results are (1) greater activation for calculation (green) in a large extent of the overall math-responsive network (bilateral IPS, IFG, anterior insula, SMA); (2) greater activation for arithmetic principles (blue) in a left-lateralized network involving angular gyrus, MFG and ventral frontal cortex (BA11) (this network is less extended for adolescents than for adults); (3) greater activation for geometry (red) primarily in bilateral inferior temporal cortex.

**Figure 12.**
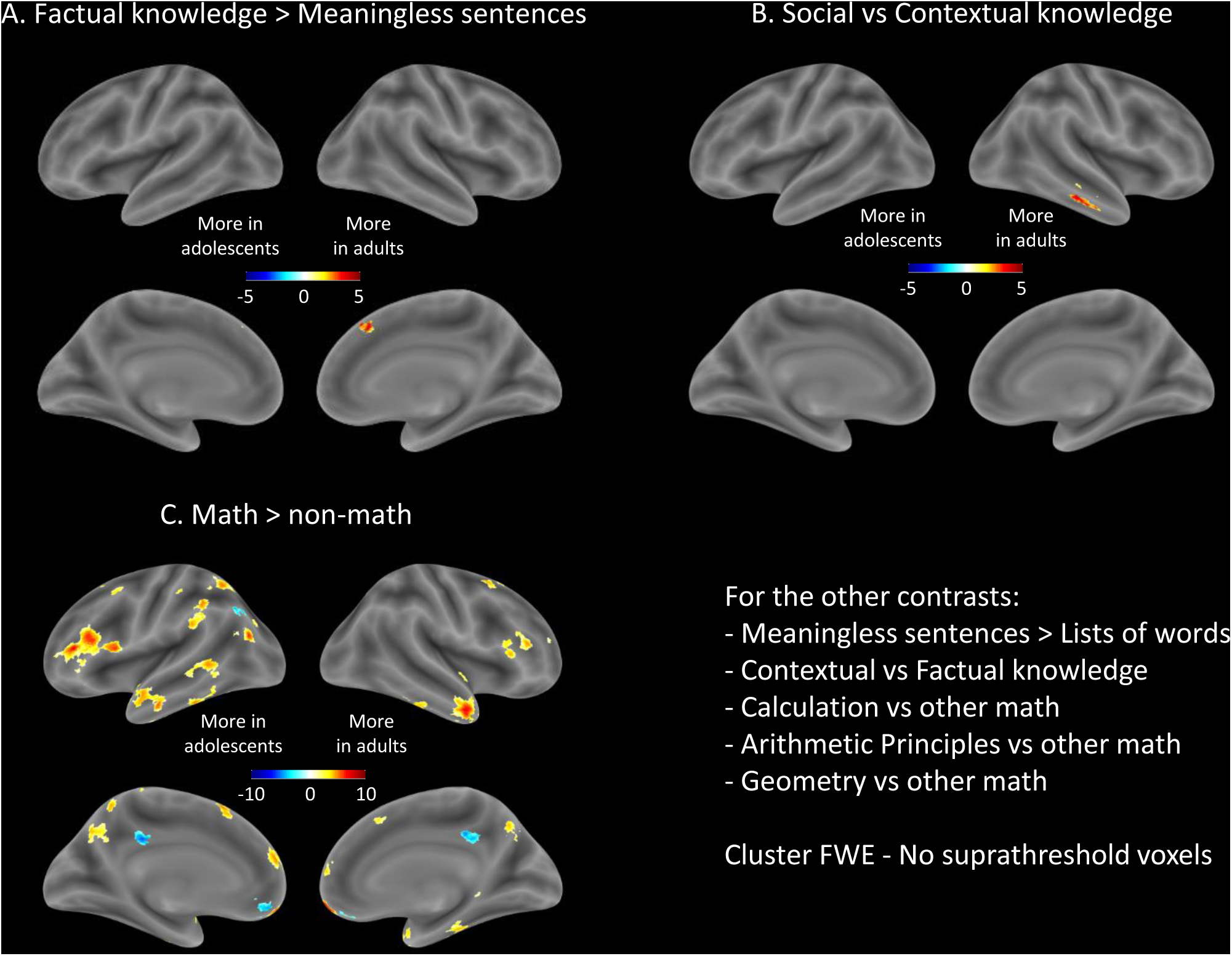
Rare contrasts showing a difference between adults and adolescents.

Another measure of similarity between adults and adolescents consisted in computing the correlation between the 16 activation levels (histograms of betas) in each group, at the same MNI coordinates. For each of the adult peaks, we found very similar response profiles and significant correlations with those in adolescents (correlation coefficients, comparing the histograms shown in figures 4-7 with those shown in figures 8-11, are presented in Supplementary materials). For instance, the highly selective response to all six conditions with mathematical statements in areas such as IPS, ITG or IFGop was almost perfectly replicated in the two groups (compare figures 6 and 10).

For completeness, figure S3 reports the interactions with modality (A or V) in adolescents. Briefly, we replicated in adolescents the findings that the left posterior ITG showed greater activation for geometry in the auditory rather than visual modality (figure S3D) and for calculation, conversely, in the visual rather than auditory modality (figure S3E). The absence of triple interactions of conditions X modality X group indicated that these subtle modality effects did not differ between adults and adolescents.

## Discussion

Previous studies established that different categories of human-specific competence, such as language, mathematics or theory-of-mind, map onto broadly different cortical networks in the human brain (Al Roumi et al., 2023; Amalric & Dehaene, 2016; Chen et al., 2021; Deen et al., 2015a; Fedorenko et al., 2011, 2012; Pinel et al., 2019; Wang et al., 2019). Our results show that it is possible, in a one-hour spoken and written sentence processing paradigm, to separate them into some of their major components. In both adults and adolescents, 3T event-related fMRI revealed the existence of distinct responses to (1) syntactic structure in sentences devoid of overall meaning; (2) elementary semantic composition of words into sentences; (3) context-dependent semantic composition taking into account a specific geographical or historical context; (4) context-specific adoption of a person’s viewpoint, an essential component of social knowledge; (5) composition of mathematical terms including numerical, arithmetic or geometric terms. We found all these networks to be amodal, responding identically to spoken and written statements. We now discuss those results in turn.

### Parsing of language areas according to syntactic and semantic contents

A classic localizer for language areas consists in contrasting meaningful sentences with meaningless lists of words (Desbordes et al., 2023; Mazoyer et al., 1993; Pallier et al., 2011) or pseudowords (Fedorenko et al., 2011, 2012). Our results show that this broad contrast can be further separated into two distinct cortical networks: one that responds to the syntactic structure of meaningless sentences, and the other that presents a supplementary response to meaningful combinations of words. As shown in figures 4 and 8, these networks are abutting in the left temporal lobe, yet syntax activates the depth of the STS, including its posterior and anterior sectors, while semantic composition yields additional activation in the middle temporal gyrus and at the posterior end of the STS, encroaching into the angular gyrus. Furthermore, within the inferior frontal gyrus, the pars triangularis and orbitalis already respond to the mere presence of syntactic structure, while semantic structure yields more anterior and dorsal activation in left prefrontal cortex. Semantic integration also involves the retrosplenial cortex. All of these cortical activations are left-lateralized, with a corresponding dissociation in two distinct sectors of the right cerebellum Crus2.

The dissociation between syntax and semantics remains a controversial topic, with a long history of early findings (e.g. Ben-Shachar et al., 2004; Mazoyer et al., 1993; Moro et al., 2001; Stromswold et al., 1996), including results (e.g. Dapretto & Bookheimer, 1999) that were later non-replicated or only partially replicated (Fedorenko et al., 2020; Shain et al., 2024; Siegelman et al., 2019). As a result, two recent articles argue that “syntactic/combinatorial processing is not separable from lexico-semantic processing at the level of brain regions” (Fedorenko et al., 2020) and that, instead, the human brain is characterized by a “distributed sensitivity to both linguistic structure and meaning throughout a broad frontotemporal brain network” (Shain et al., 2024). Our results argue against this conclusion and concur with many others in suggesting that a specific temporo-frontal network, focused on left pSTS and IFG (pars triangularis and orbitalis), plays a stronger role in syntactic than in semantic compositionality (Goucha & Friederici, 2015; Matchin et al., 2017; Pallier et al., 2011; Tyler et al., 2010; Woolnough et al., 2023), predicts syntactic performance in brain-lesioned patients (Tyler et al., 2011), and track syntactic parameters such as the number of open nodes (Giglio et al., 2024; Nelson et al., 2017).

Our results suggest that the conclusions of the Fedorenko group (Fedorenko et al., 2020; Shain et al., 2024) could be due to a methodology of first extracting voxels from the broad contrast of meaningful sentences versus lists of words or pseudowords, and then pooling together all responsive subject-specific voxels within a large region-of-interest (Fedorenko et al., 2010). This method would mix the responses of voxels which, in our paradigm, exhibit dissociable responses. The use of an intermediate condition, here semantically anomalous or “meaningless” sentences, shows that, even in the vicinity of the STS, two different types of voxels can be abutting (figure 4). We suggest that distinguishing them is of high theoretical importance: those that respond strongly to meaningless sentences, at the same level as meaningful ones, may respond to syntax, while those that respond weakly or not at all to meaningless sentences, at the same level as word lists, seem to require compositional meaning to be activated. Our group results indicate that those voxels are partially separable at the mesoscopic scale of a group 3T fMRI study, and other results by our group using single-subject precision fMRI at 7 Tesla suggest that they can also be separated in individual subjects, often in very close positions, with e.g. dorsal STS voxels more implied in syntax and ventral STS more in semantics (Dighiero et al., in preparation).

In attempting to isolate syntactic processing in the brain, using fMRI, MEG or intracranial recordings, we and other previously used “Jabberwocky” stimuli, sentences made of pseudowords, with enough function words to maintain grammatical parsability, while removing any lexical meaning (Desbordes et al., 2023; Fedorenko et al., 2016; Goucha & Friederici, 2015; Matchin et al., 2017; Pallier et al., 2011; Woolnough et al., 2023). Here, however, we use a different syntactic contrast in which meaningful words are always present, yet organized either in unstructured lists or in meaningless sentences– a condition which is more rarely studied (Goucha & Friederici, 2015; Mazoyer et al., 1993; Tyler et al., 2010). We now see disadvantages to using Jabberwocky stimuli.

One is that pseudowords typically yield weaker activation than real words throughout the language network, leading to the conclusion that lexical and compositional processes are inseparable (Fedorenko et al., 2020). However, it should be not be surprising that there are joint effects of lexicon and syntax within the same regions (Fedorenko et al., 2020; Shain et al., 2024) because part of our lexical knowledge is syntactic and records such properties of words and morphemes as part of speech; gender and number for nouns; person, tense, number and type of arguments for verbs, etc. Removing some or all of these properties in Jabberwocky may render it difficult to parse, unless great cautions is taken to preserve enough function words and morphemic markers (Pallier et al., 2011). Conversely, random word lists may still afford some syntactic construction, if chance words collide into local phrases and/or if participants can effortfully reorder them using a mental buffer (Vagharchakian et al., 2012). Such stimulus design difficulties may also explain discrepancies across studies.

The conclusion that the left fronto-temporal sentence-responsive network does not operate as a homogeneous whole, but can be partially subdivided into semantic and syntactic components, fits with the classical linguistic view, according to which language involves at least three layers of compositionality or “unification” at the phonemic, syntactic and semantic levels, each associated with distinct yet parallel fronto-temporal networks (Hagoort, 2013). This being said, other points of convergence across studies are numerous. All provide strong evidence for a major distinction between a core language network (figure 2 in Fedorenko et al., 2024), responsive to meaningless stimuli (Shain et al., 2024) and fully coinciding with the present syntactic contrast (our figure 4A), surrounded by a broader, highly distributed semantic or “knowledge” system (our figure 4B; see e.g. Deniz et al., 2019; Huth et al., 2016), unresponsive to meaningless stimuli, and which includes the left angular gyrus (Pallier et al., 2011; Shain et al., 2024). There is also agreement that both networks are amodal and respond identically to spoken and written language, as found here (Deniz et al., 2019; Fedorenko et al., 2024). Furthermore, the present results should not necessarily be seen as rejecting the idea that syntactic versus semantic distinction “is primarily a matter of degree rather than kind” (Shain et al., 2024). While fMRI primarily relies on categorical contrasts, more precise neurophysiological recordings often reveal mixtures of neuronal populations within the same region (Leonard et al., 2024) or even overlapping neural subspaces within the same neurons (Xie et al., 2022). We would not be surprised if such mixed neural selectivity for syntactic and semantic computations was ultimately found at the single-neuron level, with the present fMRI results resulting only from their different proportions across areas (see also Caucheteux et al., 2021; Pasquiou et al., 2023; Reddy & Wehbe, 2021). Indeed, while the activation to meaningless sentences sides either with meaningful sentences (syntax network) or with word lists (semantic network), some regions showed a gradual response from wordlist to meaningless and meaningful conditions, suggesting a functional overlap. Such is the case in the Cerebellum Crus2 (figure 2A and 2B): this region is deactivated by word lists and, to a lesser extent, meaningless sentences, but activates to various meaningful sentences, with a preference for social ones (figure 5B), confirming its role at a conceptual level. Our results thus confirm the important contribution of a localized subpart of the cerebellum to language even in the absence of overt production (Casto et al., 2025; LeBel et al., 2021; Schmahmann et al., 2019).

### Cortical specialization for contextual and social semantic processing

Among the brain networks engaged in semantic processing, a novel finding of the present work is that the requirement to modulate word meaning by local context leads to a sudden and non-lateralized increase in activation in a well-delimited set of bilateral regions: the angular gyri and temporal poles. This is consistent with previously findings of semantic integration in both regions (Bemis & Pylkkänen, 2013; Bonnici et al., 2016; Mazoyer et al., 1993; Pallier et al., 2011; Price et al., 2015, 2016). However, the present short-sentence paradigm introduces a novel minimal contrast to engage them, showing that they can be selectively triggered by the mere introduction of a few contextual words such as “In London” or “For Einstein”, thus requiring to interpret subsequent words within this semantic background. It is striking that this manipulation has a strong impact on a selective subset of brain regions (figure 5A), while leaving other syntax and semantic-related areas essentially unchanged (see the histograms in figure 4B). Vector-based models of semantic composition suggest that individual words are encoded as vectors in conceptual space (Landauer & Dumais, 1997; Pennington et al., 2014) whose meanings can be combined by operations of vector addition and combination, possibly explaining the ramping brain signals observed when successive words get integrated into phrases (Desbordes et al., 2023; Nelson et al., 2017; Pallier et al., 2011; Woolnough et al., 2023). While the angular gyrus is specifically involved in such combinatorial semantics, even for two-word combinations (e.g. “plaid jacket”; Price et al., 2015, 2016), computational models indicate that non-linear operations are sometimes needed to combine word meanings (Baroni & Zamparelli, 2010; Socher et al., 2012)—for instance the meaning of “red guard” is not the sum of its component words. Our contextual sentences put a particular emphasis on such transformations, because they were designed such that context could reverse the truth value of the embedded phrase. Our results therefore hint at a possible role of bilateral AG and TP in the non-linear vector operations that contextualize semantic embeddings. This conclusion is compatible with the many observations that the activation of these regions are well captured by mid-layer embeddings of large-language models, which implement such transformations, over and above the addition of single-word embeddings (Caucheteux et al., 2022; Pasquiou et al., 2023; Schrimpf et al., 2021).

The vast majority of sentences in our “contextual” condition referred to geographic contents (e.g. “in London…”), and this may explain some of the observed dissociations. While both geographic and social contexts jointly activated the bilateral angular gyrus and temporal poles, other areas showed domain-specificity. The left and right mesial retrosplenial and parahippocampal cortices were selective to geographic context (blue in figure 5B), fully compatible with much previous evidence for their role in spatial cognition (Mitchell et al., 2018; Vann et al., 2009; Woolnough et al., 2020), responding not only during spatial orientation or picture processing, but also story listening (Huth et al., 2016) and access to large-scale geographic knowledge (Peer et al., 2019). Conversely, the right TPJ was quite selectively activated by social context (figure 5, bottom), compatible with its previous involvement in theory of mind (Deen et al., 2015a; Saxe & Kanwisher, 2003). The double dissociation was most evident in the midline posterior cortex, where two adjacent regions responded to social (bilateral precuneus) versus contextual knowledge (retrosplenial cortex), respectively (figure 5B). In addition, social-context judgements caused an extended overactivation of a broader but less specific left-lateralized language network (figure 5), all along the left STS, in agreement with prior work (Deen et al., 2015b, 2023; Deniz et al., 2019; Huth et al., 2016; Mar, 2011). The rich semantics of familiar person names, evoking historical, scientific, professional and emotional knowledge, probably contributed to this extended activation (Desai et al., 2023). Note that our social and contextual sentences contained roughly matched numbers of proper names (39 vs 35), but our results are compatible with observations that persons, more than places, activate bilateral anterior and posterior temporal regions (Desai et al., 2023; Huth et al., 2016). The overlap of brain networks for mentalizing (thinking about other people’s thoughts) and for sentence and story comprehension has been noted by others (Huth et al., 2016; Mar, 2011), but dissociations are also on record within the STS (Deen et al., 2015b), and more work at the single-subject level is needed to fully resolve this issue.

### A distinct amodal math-responsive network

Our findings support the existence of a distinct math-responsive brain network, radically different from language areas, that processes a variety of math concepts. This observation replicates and extends prior work from our laboratory in mathematicians (Amalric & Dehaene, 2016, 2019). Here, participants were not mathematicians, but schooled adults (figure 6) and adolescents (figure 10), yet we still observed robust activation to mathematical statements in a bilateral network including intraparietal sulci (IPS), inferior temporal gyri (ITG) and middle frontal gyri (MFG) – a classic set of regions previously found during mental arithmetic (Dehaene et al., 2003; Pinheiro-Chagas et al., 2018, 2024), but also more recently in geometry, number line and graphicacy tasks (Aulet et al., 2025; Ciccione & Dehaene, 2025; Dehaene et al., 2025; Sablé-Meyer et al., 2025). This network was remarkably similar between adolescents and adults, the only difference being a greater left prefrontal cortex activation in adults, particularly in the left MFG, which could reflect greater cognitive effort due to the long interval since formal math training. In both age groups, the double dissociation with the semantic network evoked by general-knowledge sentences was radical, with one being deactivated below the resting-state baseline in exactly the conditions where the other activated (histograms in figures 6 and 10). In agreement with others (Cantlon et al., 2006; Morfoisse et al., 2025; Nakai et al., 2023), our results thus suggest that a distinct math-responsive network exists independently of age and math expertise.

Importantly, in both age groups, the math-responsive network was essentially independent of presentation modality. This amodal selectivity aligns with previous findings that numerical processing, particularly in the IPS, is independent of whether numbers are presented visually or auditorily (Eger et al., 2003), and is consistent with evidence that even individuals blind from birth engage the same core math network, including IPS, ITG and MTG (Amalric et al., 2018; Kanjlia et al., 2016). Here, such amodal responses also generalized across different math domains, including arithmetic principles, calculation, and geometry. The posterior ITG activation to geometry did exhibit an effect of modality, with greater activation for auditory than visual presentation, possibly indicating interference from visual sentences onto the mental images evoked by geometric statements. However, this small effect should not detract from the broader finding that the core math-responsive network, including a more anterior portion of the ITG, responded robustly to all math statements across both modalities.

The present findings address several potential objections or confounds that might have affected previous observations of a math-responsive network (Amalric & Dehaene, 2017). One common argument is that math cognition is primarily spatial in nature, which might explain its reliance on parietal areas. However, while spatial reasoning likely contributes to geometric thinking, and numbers and calculation evoke a mental number line (Knops et al., 2009), it is much less clear why knowledge of formal arithmetic principles (e.g. multiplication by zero) would also rely on spatial maps, and previous work also found the same network active during verification of rote algebraic expressions (Amalric & Dehaene, 2019). Another concern is whether our findings could be explained by intrinsic task difficulty. However, math statements were not systematically more difficult than non-math statements, in fact calculation was the least error-prone condition (figures 2 and 3).

Moreover, the core math network remains selective for math statements, even when compared to equally or more demanding non-math conditions (present results, and Amalric & Dehaene, 2016). Taken together, our findings reinforce the idea that rather than mere effects of spatial processing, difficulty or an ill-defined domain-general “multiple-demand” system (Duncan, 2010), the observed activation reflects the operation of a network sensitive to various mathematical concepts and patterns.

### Further specialization within mathematics

Our results also reveal finer-grained specialization within mathematics. Calculation caused a greater engagement of a fronto-parietal circuit encompassing the horizontal segments of the IPS, precentral and inferior frontal regions, supplementary motor area, and the anterior insula. This pattern is consistent with previous findings linking these regions to procedural and rule-based operations and working memory (Arsalidou & Taylor, 2011; Jost et al., 2009). The relative increase in left prefrontal activity during calculation might therefore reflect an increased demand on maintenance and manipulation of intermediate results in working memory. By contrast, sentences expressing arithmetic principles elicited greater activation in left middle and inferior frontal and angular gyri, as well as in the left middle temporal gyrus, suggesting that these regions are involved in the acquisition of abstract, relational knowledge, as previously shown during abstract logical inference (Monti et al., 2009). Interestingly, some of these regions also showed an increased activation to social-knowledge statements (figure 7, bottom); whether the same neural systems support abstract reasoning and rule integration across the math and social domains is an intriguing suggestion that merits further investigation.

Finally, geometry, relative to other math statements, produced a distinctive bilateral activation in posterior ITG, replicating other findings from our lab (Amalric & Dehaene, 2016; Morfoisse et al., 2025). These regions lie just anterior to the occipito-temporal boundary and form the posterior tip of the broader inferior temporal activation observed across all math statements, suggesting a possible gradation of math knowledge along the posterior-anterior axis of the ITG, from more visual to more abstract representations (see also Popham et al., 2021). The posterior localization and stronger response to visually presented sentences are consistent with the idea that geometrical thinking elicits additional visual imagery or shape-based processing, in line with prior research on mental imagery (Liu et al., 2025). Indeed, in our previous study of professional mathematicians (Amalric & Dehaene, 2016), self-reported mental imagery during reflection on spoken math statements correlated with activity in the left posterior ITG.

Thus, while a core math-responsive network appears amodal, our results suggest that different mathematical domains recruit partially distinct subnetworks, depending on the representational format and cognitive demands involved. Geometry may rely more heavily on visuospatial and imagery-based processes, arithmetic principles on abstract rule integration, and calculation on procedural and working memory mechanisms.

### Methodological aspects and limitations

The present design allows to efficiently parse various regions of the language and math-response networks. A limit of the present analyses is that they are performed at the group level, but ongoing work in our laboratory suggests that the findings replicate at the single-sentence level (Pietrantoni et al, in preparation) and in single voxels of single subjects using 7T fMRI (Dighiero et al., in preparation). In this respect, it is quite similar to other efforts to obtain efficient fMRI localizers separating language, social and math networks (Hutchinson et al., 2024; Pinel et al., 2007; Pinho et al., 2018; Tuckute et al., 2024). However, unlike (Tuckute et al., 2024), we do not consider it sufficient to contrast meaningful sentences versus list of pseudowords, and then pool together all active voxels within relatively large ROIs (Fedorenko et al., 2010). The present work clearly suggests that finer anatomo-functional distinctions can be achieved by using intermediate stimuli, including lists of words (carefully crafted to avoid creating syntactic constituents) and meaningless sentences (carefully crafted to permit syntactic parsing while minimizing semantic content).

Another limitation of the present work is that stimulus design did not necessarily permit a full separation of the relevant cognitive levels. Most crucially, although the semantically anomalous sentences were called “meaningless” for simplicity, their syntax afforded thematic assignments (“who does what to whom”) and their beginnings were often meaningful, thus potentially causing transient semantic activations. For lack of time, we also did not include control conditions such as lists of math words or meaningless math statements—but we did in previous work (Amalric & Dehaene, 2016), with very similar results indicating that the math-responsive network is sensitive to meaningful composition.

### Conclusion

While natural language, mathematics and social cognition all involve the composition of words into higher-level syntactic and semantic structures, the present results confirm that those domains rely on partially dissociable parallel circuits temporo-parieto-frontal circuits (Dehaene et al., 2022). Even within language, partially different subsystems are involved in syntactic and semantic compositionality (Hagoort, 2013). In addition to involving parallel cortical circuits, most of these networks also comprised a cerebellar component, in agreement with much recent research suggesting that distinct parts of the cerebellum are involved in high-level cognition, including semantic-level operations in math and social domains (Buckner et al., 2011; Guell & Schmahmann, 2020; LeBel et al., 2021; Schmahmann et al., 2019). The present paradigm can serve as a localizer to reliably identify them, possibly using a fraction of the present stimuli, and then further dissect their response properties in single subjects, arguably the best practice in fMRI (Fedorenko et al., 2010; Shain et al., 2024). In particular, now that the present work conclusively demonstrated that these activations are amodal, future work could rely on a single modality, for instance written sentences for high-field MRI studies where presenting auditory stimuli can be difficult, or spoken sentences for blind participants (Amalric et al., 2018; Kanjlia et al., 2016).

## Acknowledgements

This research was supported by INSERM, CEA, Collège de France, Université Paris-Saclay, the Bettencourt-Schueller foundation, the Agence Nationale de la Recherche France 2030 program, under reference ANR-23-IAHU-0010, ERC grants “NeuroSyntax” and “MathBrain” to S.D., and the European Union (FEDER-2007–2013, agreement #91–2015–004) supporting the 3T Prisma magnet at NeuroSpin. We are grateful to all support cells at NeuroSpin for their help with subject recruitment, scanning and data processing, and to the CMRR lab in Minneapolis for their multiband EPI sequence.

## Supplementary figures

**Figure S1.**
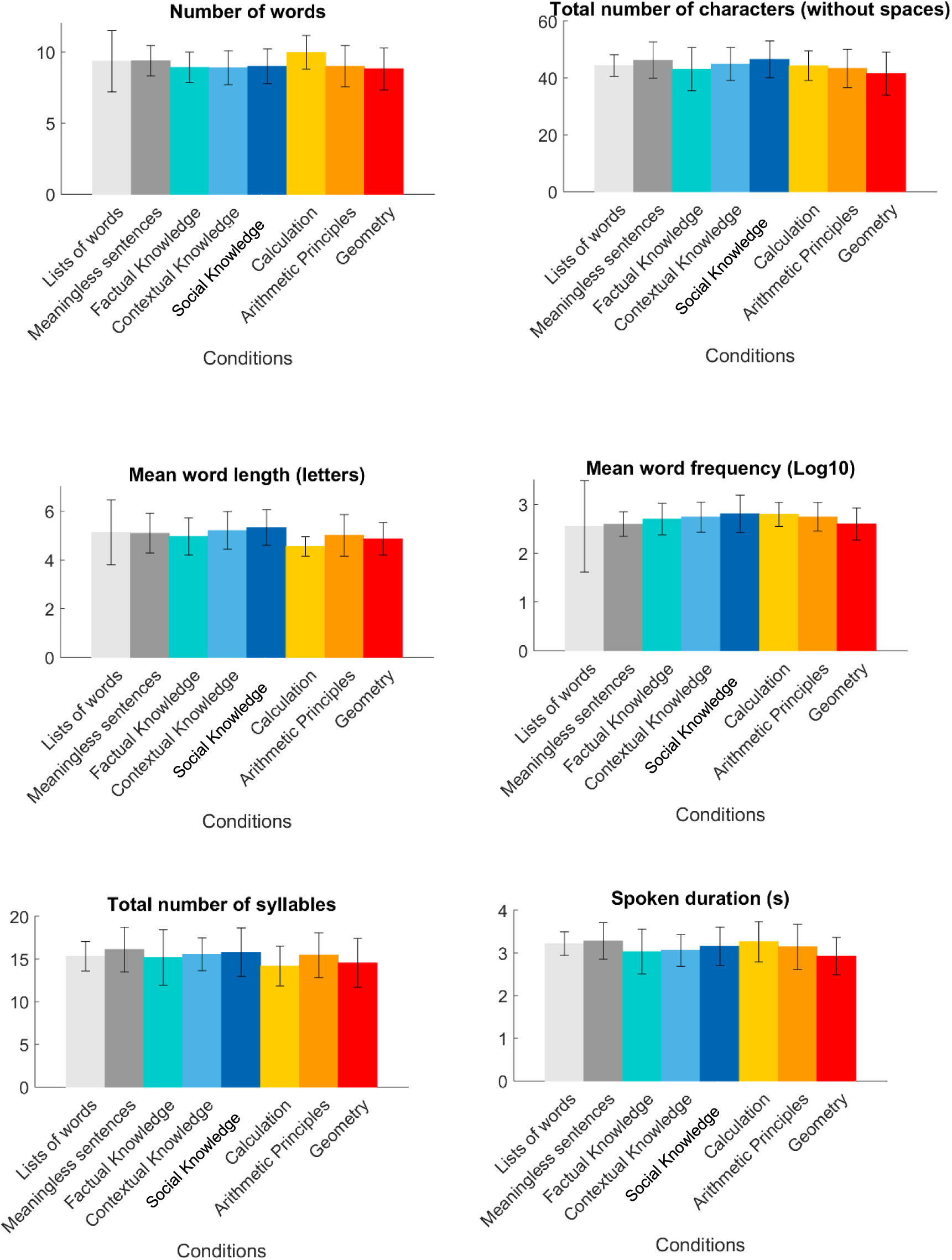
Comparison of the 8 categories of statements along six different stimulus dimensions. Each bar indicates the mean +/- one standard error.

**Figure S2.**
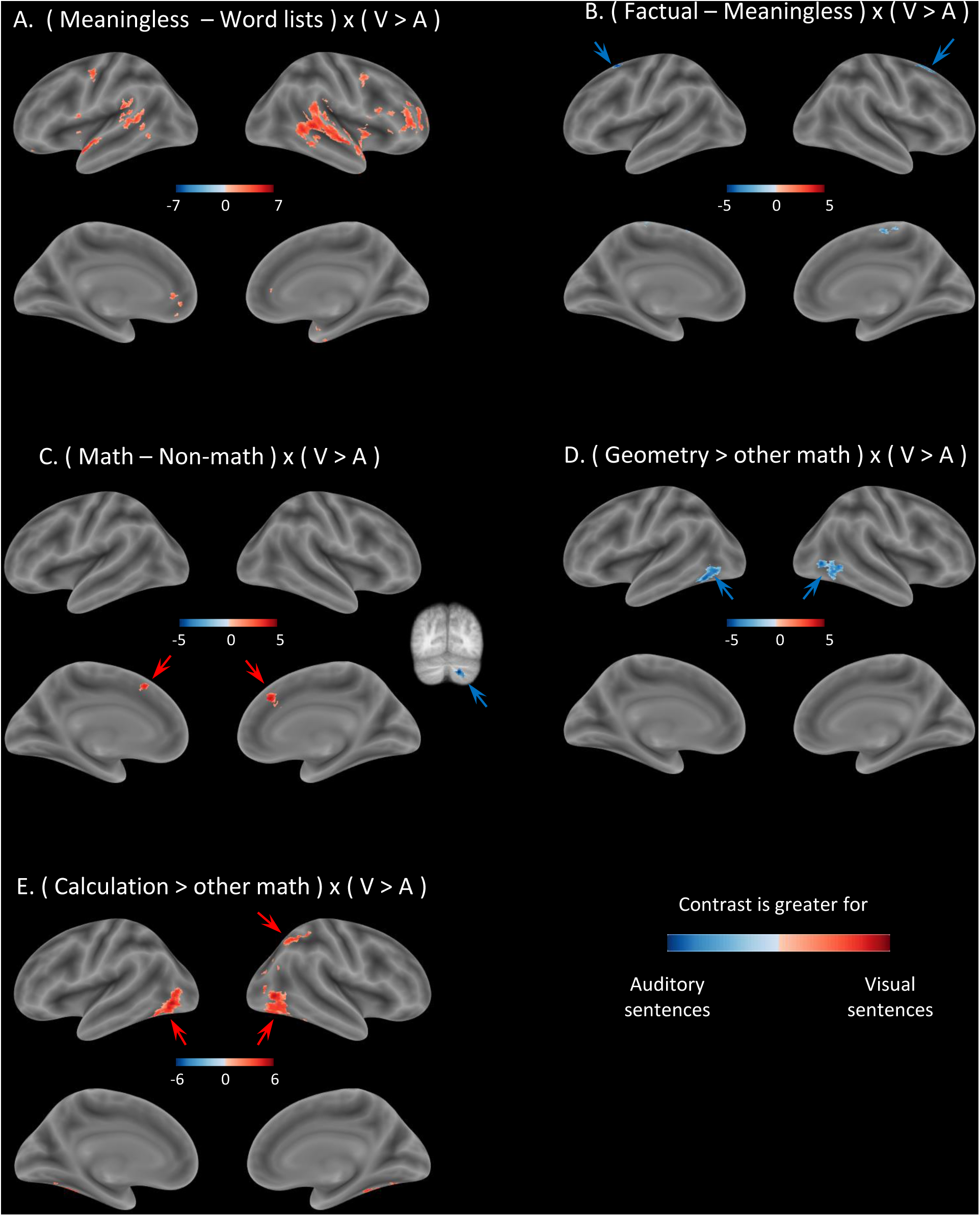
Rare interactions of modality with the main contrasts in adults. Each panel reports the interaction of the indicated contrast with modality contrasts (V>A). All images are thresholded at voxelwise p<0.001 and FWE-corrected cluster-level p<0.05).

**Figure S3.**
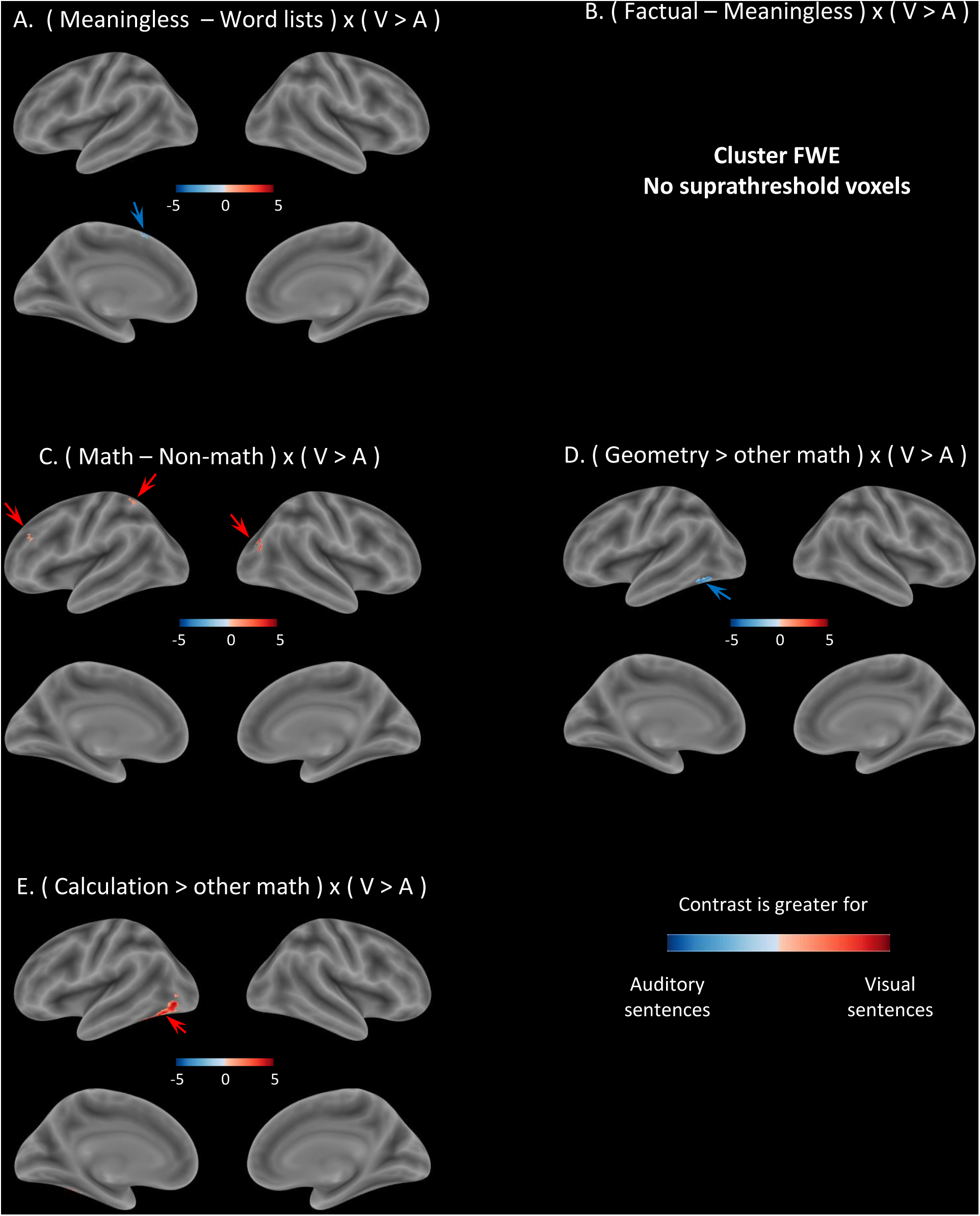
Rare interactions of modality with the main contrasts in adolescents. Each panel reports the interaction of the indicated contrast with modality contrasts (V>A). All images are thresholded at voxelwise p<0.001 and FWE-corrected cluster-level p<0.05).

## Supplementary material

**Tables of coordinates for the main fMRI contrasts**

Tables of the fMRI contrasts:

**Table S1.**
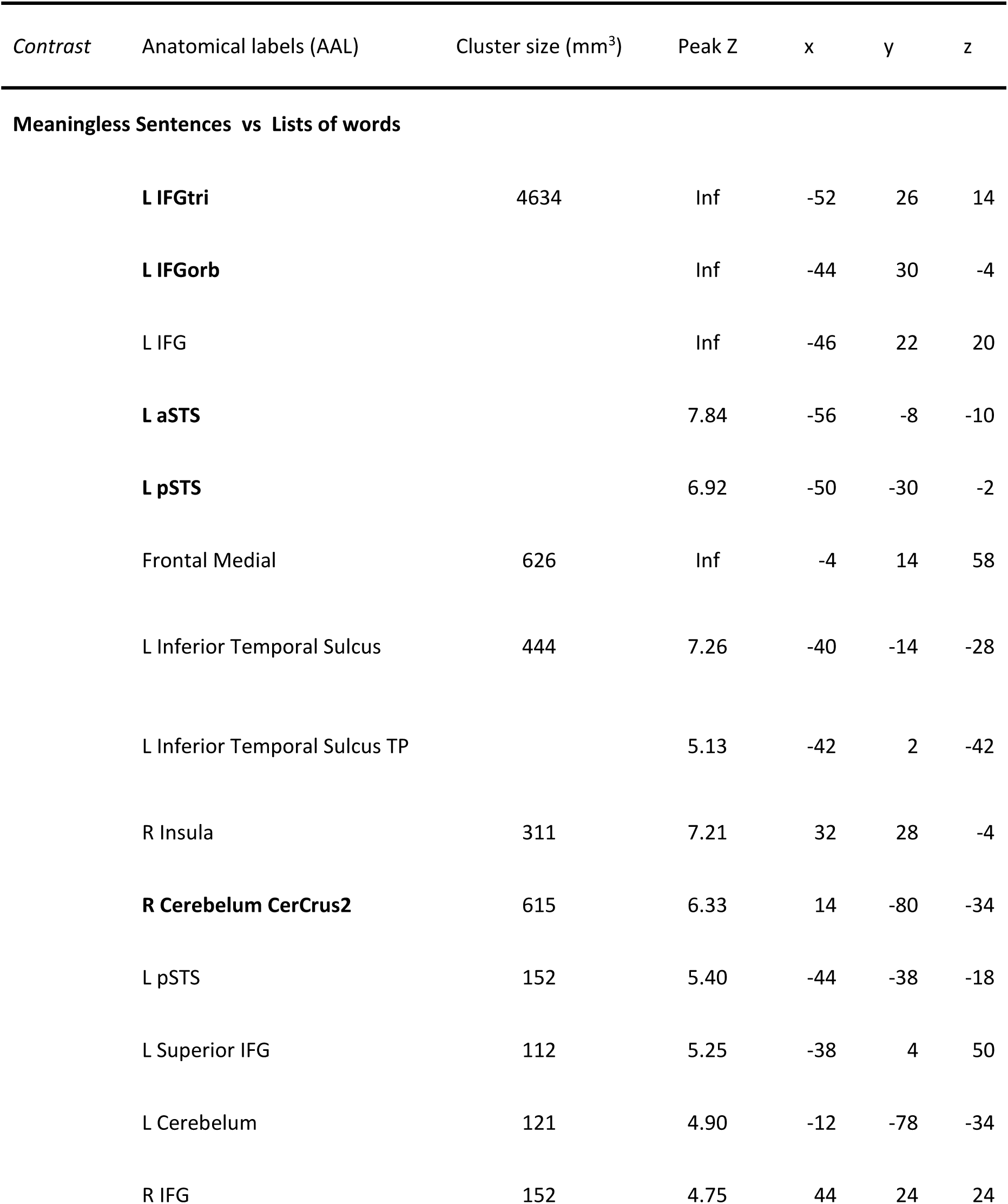
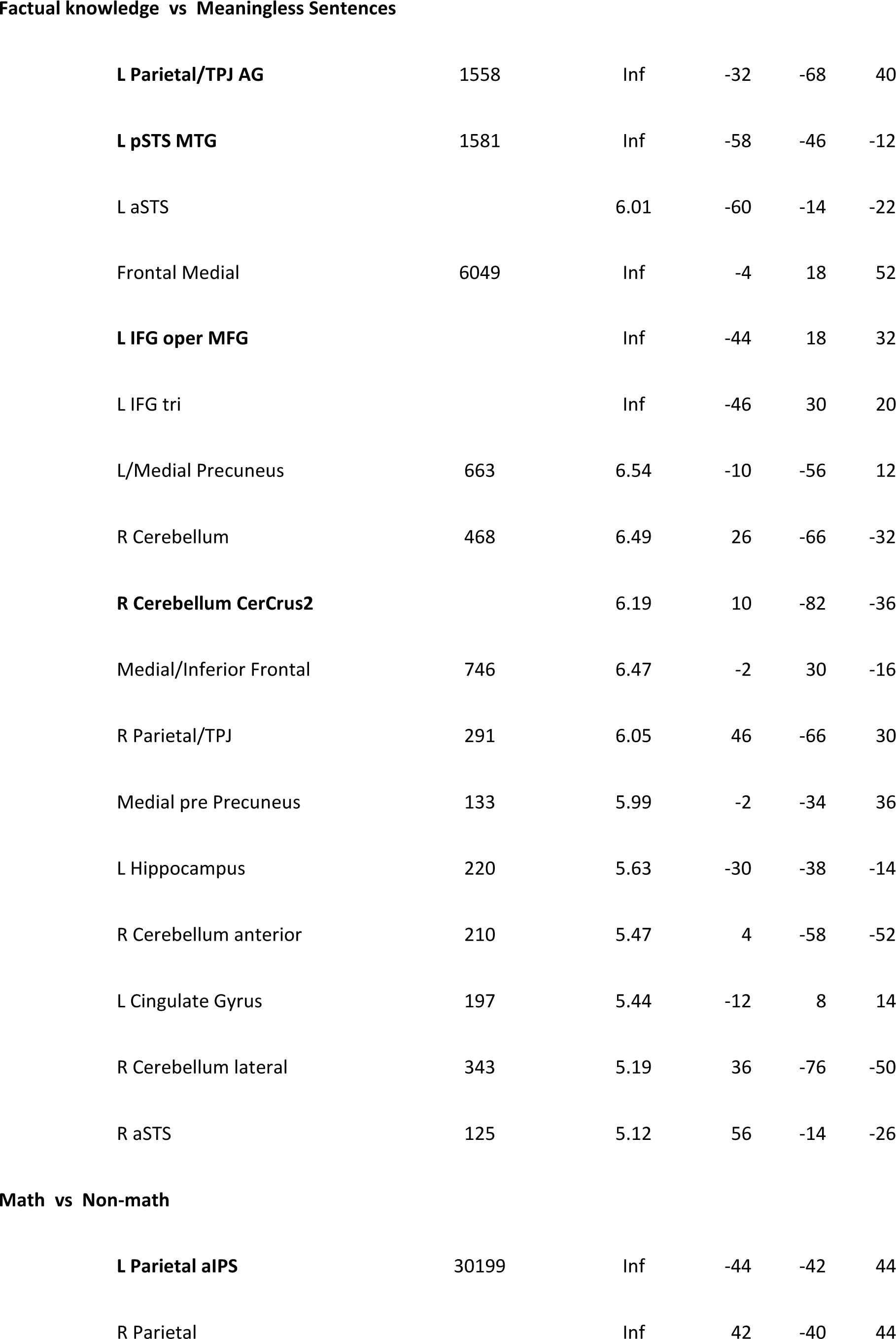
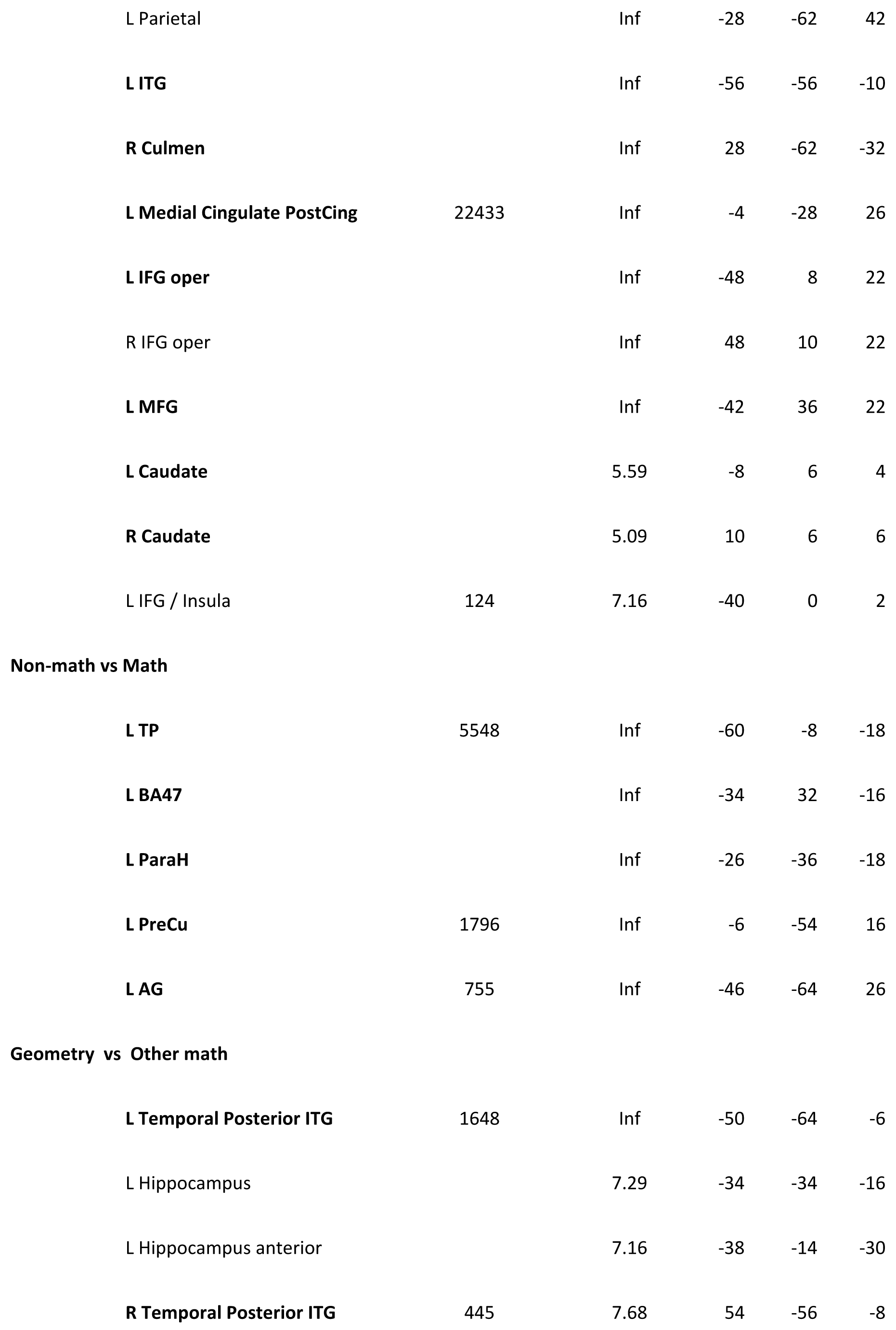
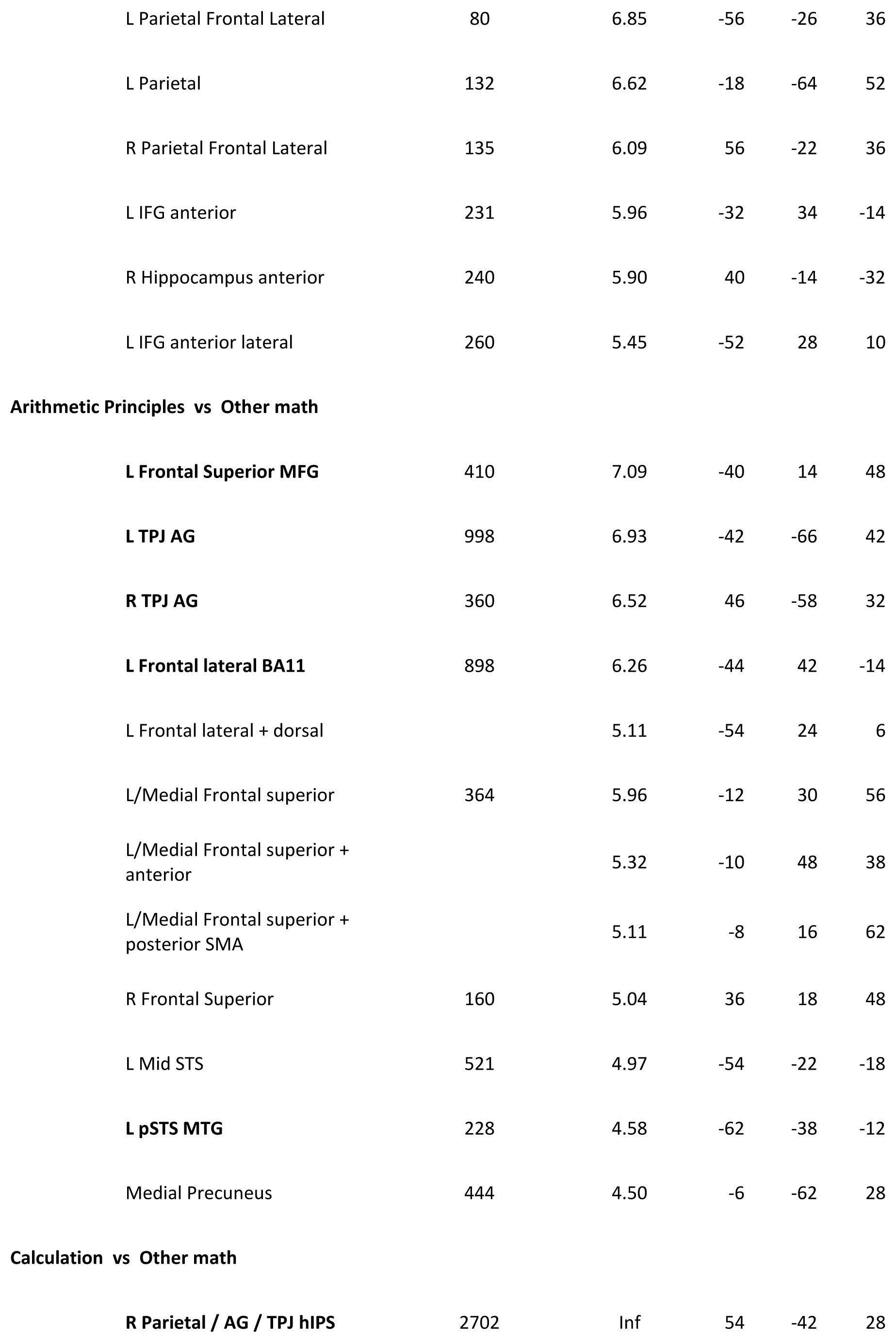
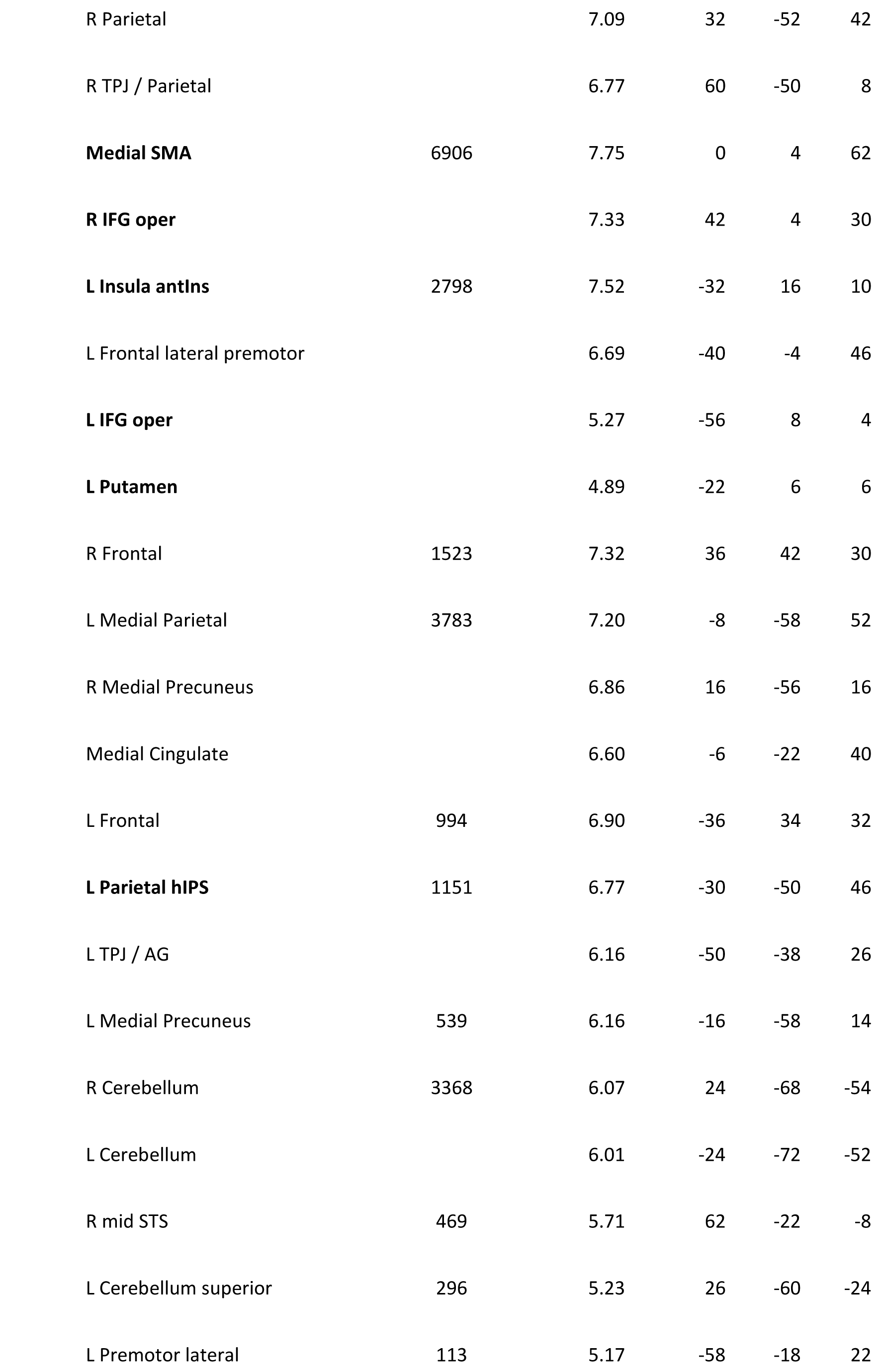
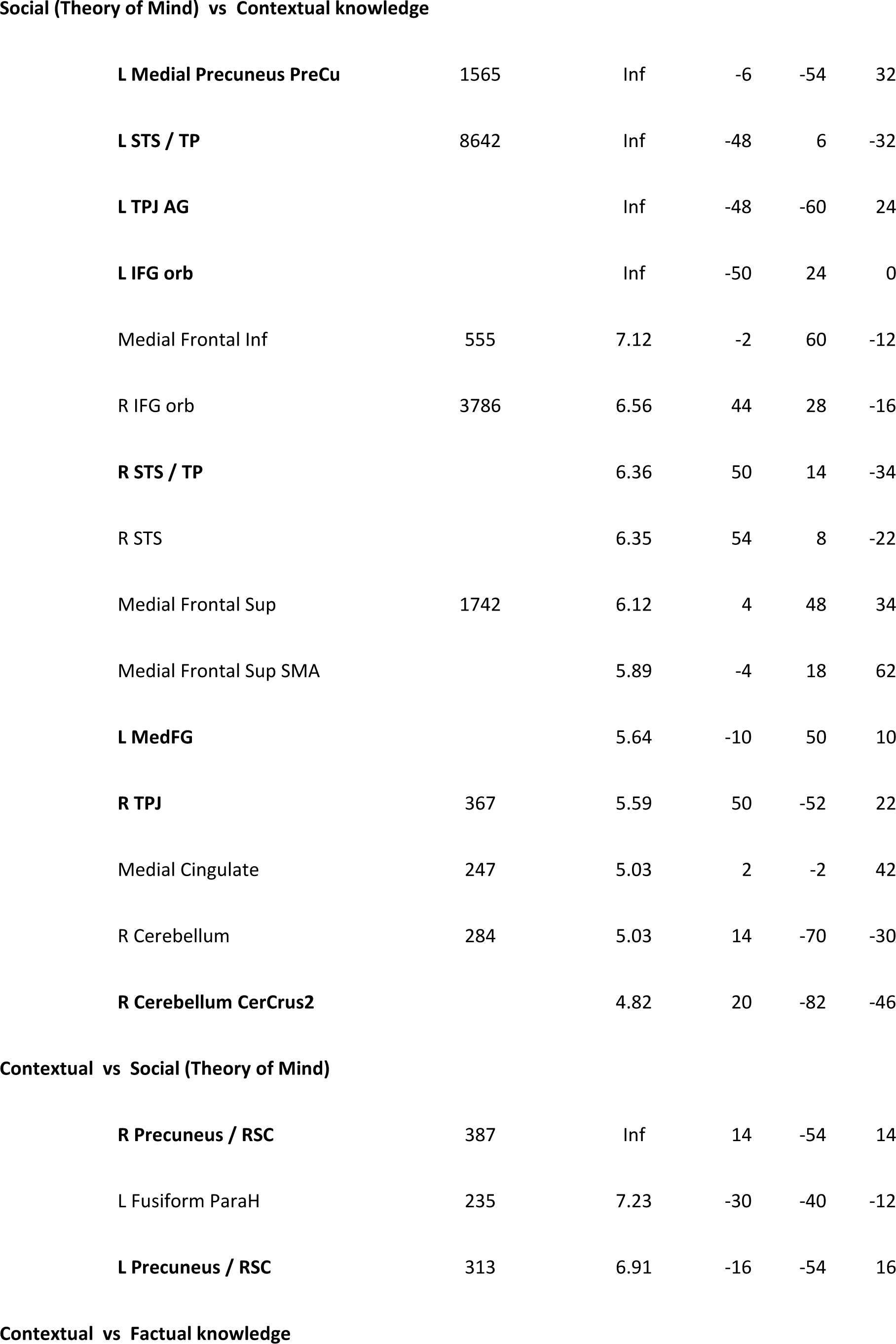
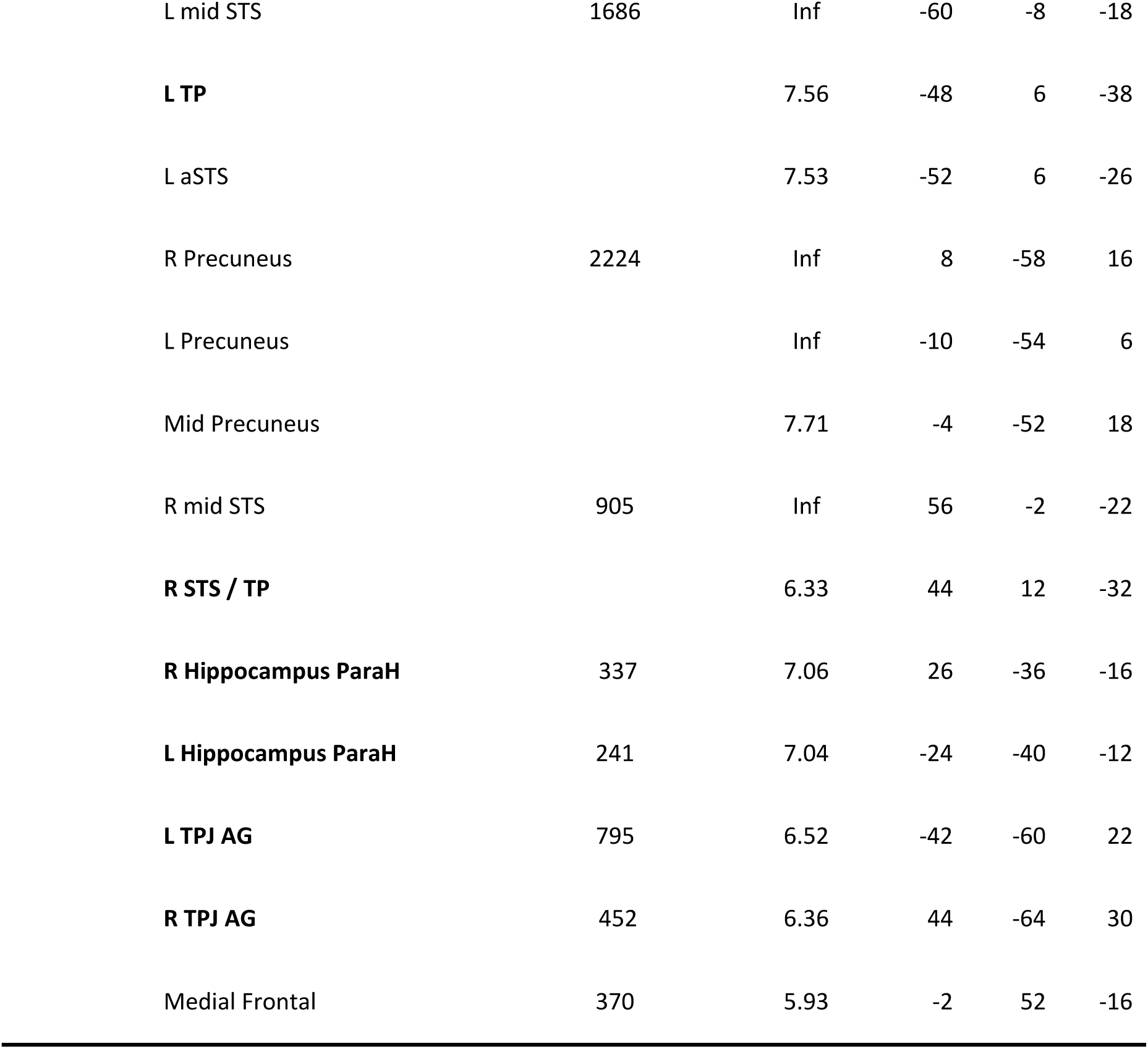
Significant contrast peaks for the different contrasts. Coordinates are in MNI space. A p=0.001 voxelwise threshold, uncorrected for multiple comparisons, was applied. Anatomical labels were obtained with the Anatomical Automatic Labeling toolbox (Tzourio-Mazoyer *et al*., 2002) (AAL version: aal3v1). In **bold** are the peaks appearing in figures 4 to 7.

**Correlation of activation profiles of adults and adolescents**

To assess intergroup reliability, a Pearson correlation coefficient R between the adult and adolescent beta was computed across the 16 conditions of the experiment, at the peak sites derived solely for adult data shown in figures 4-7.

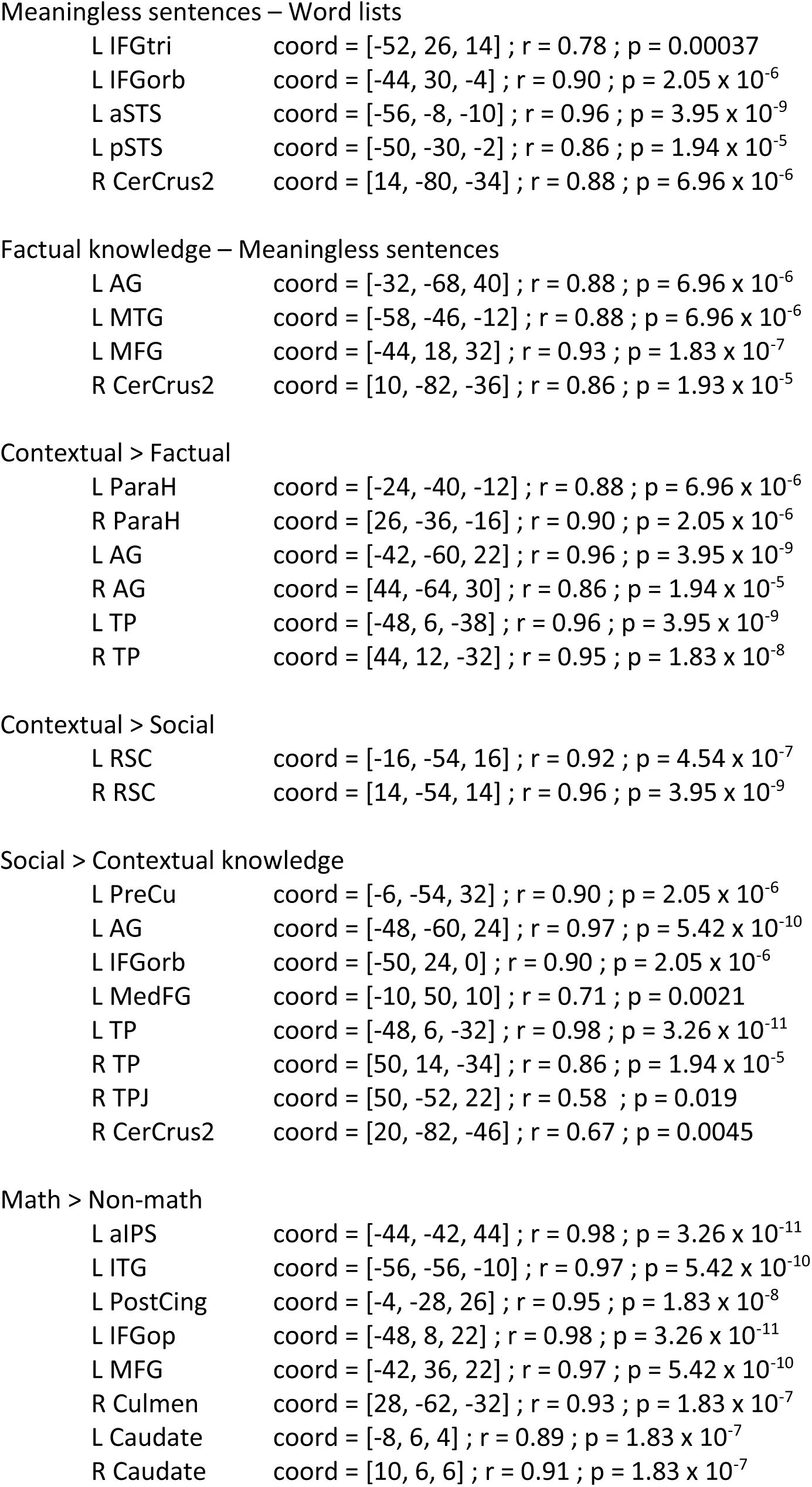
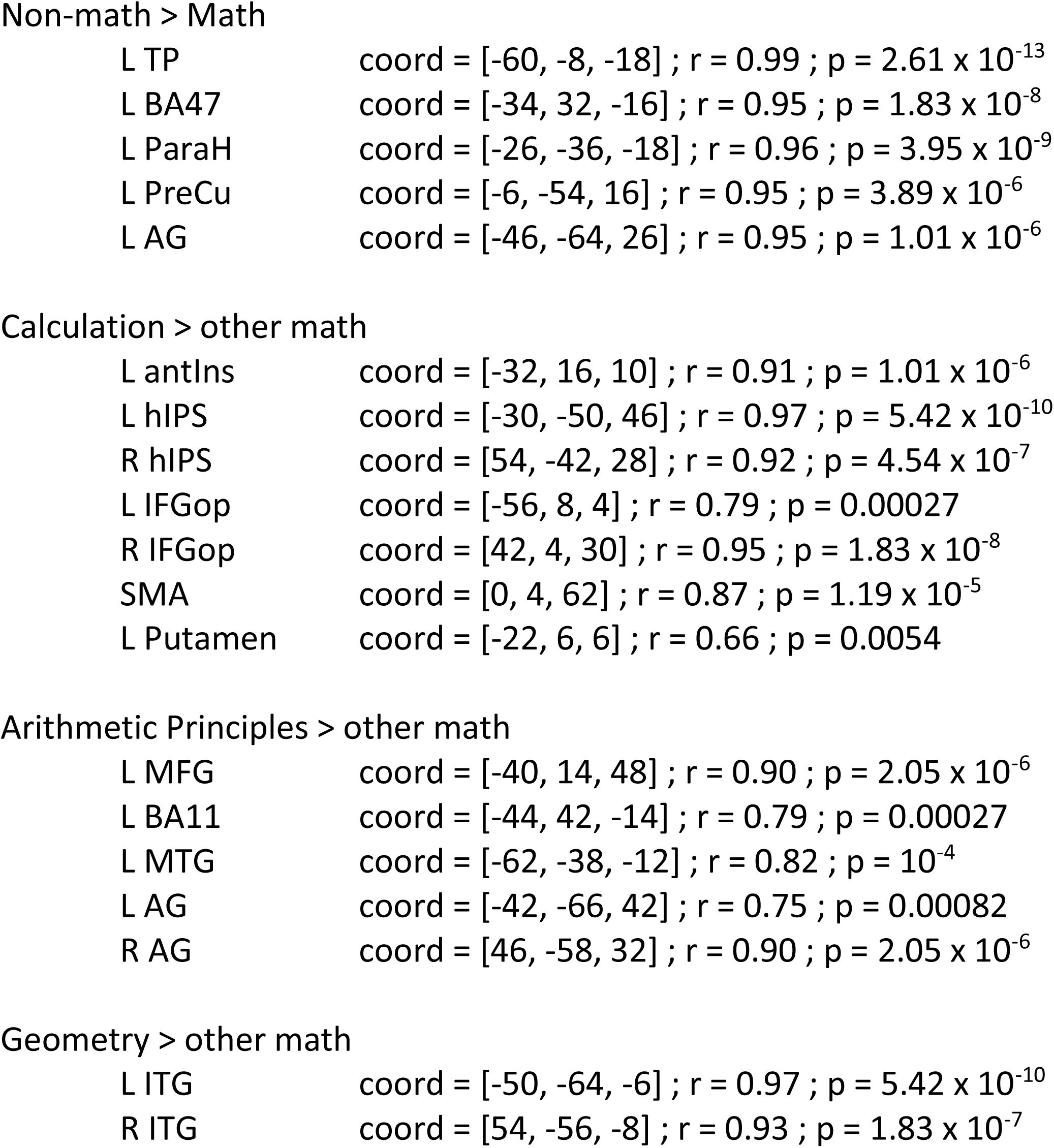

